# Simultaneous *in vivo* Pharmacokinetics and Pharmacodynamics of TDF Intravaginal Ring during High-dose Vaginal SHIV Challenge in Pigtail Macaques

**DOI:** 10.1101/2021.06.29.450465

**Authors:** Katarina Kotnik Halavaty, Adina K. Ott, Danijela Maric, Jonathan T. Su, Edgar Matias, Edward J. Allen, Amy Martin, Lara Pereira, Frank Deyounks, James M. Smith, Patrick F. Kiser, Thomas J. Hope

## Abstract

The demonstration of complete protection of macaques in a repeated low dose virus challenge by a tenofovir disoproxil fumarate (TDF) intravaginal ring (IVR) and the success of the dapivirine IVR in clinical trials highlighted the potential of IVRs as pre- exposure prophylaxis against HIV. Efficacy of TDF ring was not investigated in sexually active women. Our understanding of the mechanisms of protection is limited. To address this knowledge gap, we performed simultaneous pharmacokinetic and pharmacodynamic analysis of a TDF-IVR at the site of SIV challenge in pigtail macaques at the anatomical and cellular level. Specifically, we challenged TDF-IVR administered pigtail macaques with a single high dose of a non-replicative SIV-based vector containing a dual reporter system that helped us to identify the earliest targets of SIV infection within the mucosa. Two and three days after challenge, the macaques were euthanized and tenofovir (TFV) concentrations were measured in the female reproductive tract (FRT) by HPLC-MS/MS to correlate drug concentrations and SIV-vector transduction efficiency. TFV formed a gradient through the mucosal tissue, with the highest concentrations near the ring, in the upper vagina and endocervix. Despite this, several transduction events were identified with the most common sites being in the ovaries. Moreover, proviral DNA was detected in the cervix and vagina. Thus, our studies demonstrate an uneven distribution of TFV in the FRT of macaques after release from a TDF-IVR that leads to incomplete FRT protection from high viral dose challenge.

## Introduction

Over the past decade, antiretroviral (ARV) HIV prevention strategies have undergone numerous clinical failures and some remarkable successes in the use of oral Truvada, oral Descovy (1–4) and injectable cabotegravir. Yet, most of what we know about ARV prevention of sexual transmission through the female reproductive tract (FRT) has been learned from clinical studies, where the main outcome is the correlation of local drug levels with reduced rates of infection among women (5–9). The CAPRISA- 004 clinical trial results revealed that 1% tenofovir (TFV) gel applied vaginally before and after coitus reduced the risk of HIV acquisition among women in South Africa by 39% (5). Despite encouraging results, a subsequent VOICE study resulted in ineffective protection of coitus-independent daily use of this 1% TFV gel in female participants (6). Similarly, no evidence for efficacy in HIV prevention was found by researchers who conducted the FACTS 001 Phase III trial (9, 10). In a secondary analysis of the FACTS trial data, detection of TFV in vaginal fluids was significantly associated with a 52% reduction in HIV acquisition (9). Furthermore, a silicone-elastomer matrix IVR eluting the non- nucleoside reverse transcriptase inhibitor (NNRTI) dapivirine (DPV) has been tested in two separate clinical trials, ASPIRE (MTN-020) and Ring Study (IPM-027) in Sub-Saharan Africa. The outcome of the ASPIRE phase IIIb clinical trial demonstrated overall protection by 37% against HIV-1 acquisition (7), which was lower than initially expected. Further analyses revealed that the ring could reduce HIV risk between 56% (7) and 75% (11) depending on consistency of the ring usage. In open-labeled settings a 39% reduction in HIV-1 infections among populations with high background of HIV-1 incidence was observed (12).

In this study we utilized a non-human primate model that enabled us to study a mechanism of protection by TDF-IVR in a controlled and sustained release manner locally. We previously showed that a 28 day TDF-IVR can prevent systemic SHIV infection in pigtailed macaques (PTM) after multiple low-dose vaginal challenges (13, 14). However, the mechanism of this protection remained unexplained. The efficacy of the TDF-IVR in preventing systemic infection could either be due to high local FRT drug concentrations blocking infection from low-dose challenge all together (sterilizing protection) or lowering the frequency of early infection events and containing viral spread in a way that impedes systemic infection and viremia. Waiting several days for the basic read out of systemic infection does not allow for the exploration of important mechanistic details on the biology and pharmacology of the prevention strategy under examination. In contrast, simultaneously investigating the viral and pharmaco-dynamics and pharmacokinetics of virus-drug interaction during the first few hours post-challenge leads to a better understanding of the mechanisms through which prevention strategies may interfere with viral dissemination in the complex environment of the FRT. The two main points to be addressed to understand these mechanisms are the biodistribution of drug in the mucosal site of challenge and the sufficiency of local drug concentrations to prevent all infection throughout the entire FRT in a high-dose challenge model.

A Phase 1 clinical study showed that TDF-IVR was safe, well accepted and well tolerated with protective TDF levels in sexually abstinent women over 14 days of continuous ring use (15). However, the following Phase 1b study was terminated early due to the development of genital ulcers near the ring and high levels of multiple inflammatory cytokines and chemokines in sexually active women administered with TDF-IVR over 90 days (16). These data raise new toxicological questions concerning drug concentrations generated in the FRT with the TDF-IVR.

We have previously developed a replication defective pseudoviral dual reporter system (LICh) that allows the identification of potential foci of transduction 48 hours after exposure in the FRT (17). These foci can be further analyzed by characterizing transduced cells based on mCherry and luciferase genes expression using immunohistochemical staining. This study revealed that the entire upper and lower FRT could be exposed to the vaginal challenge inoculum and contain susceptible target cells. It was not clear before this study that the upper FRT of rhesus macaques, and particularly the ovaries were susceptible to viral transductions following vaginal challenge with a single high-dose of this pseudoviral LICh reporter (17). These observations raised questions whether an IVR would provide enough antiretroviral drug to the entire FRT to protect the upper FRT including ovaries.

In the current study, we employed this LICh reporter system to identify the locations within the FRT of early SHIV transduction in PTM protected by TDF-IVR treatment. In parallel, at necropsy, 3 days after challenge, we were able to collect the whole FRT and measure drug concentrations throughout this organ. In this way, the LICh system allowed us to run pharmacodynamic (PD) studies simultaneously with pharmacokinetics (PK) analysis at the anatomical and cellular level which revealed uneven tissue drug distribution and random viral transduction within the PMT FRTs.

## Results

### Transduced cells are present in the ovaries and lymph nodes of TDF-IVR treated macaques after a high-dose of SHIV-LICh challenge

In this study, we utilized our dual reporter LICh system pseudotyped with the HIV-JRFL envelope protein that allowed us to localize and phenotype cells transduced in untreated rhesus macaques (17) to study TDF-IVR function in our PTM vaginal challenge model. Briefly, the dual reporter LICh system carries firefly luciferase (18) and fluorescent mCherry (19) genes that are driven by a CMV promoter (Fig. S1). IRES (internal ribosome entry site) enables efficient expression of mCherry gene whereas WPRE (Woodchucks Hepatitis Virus posttranscriptional regulatory element) at the 3’ end of the genome enables increased gene expression of the vector (20, 21). The 5’ end long terminal repeat (LTR) contains the SIV promoter for efficient virus production. The 3’ end LTR site has a self-inactivating mutation. The reporter genome does not contain any other viral genes and is delivered by a SIV-based lentiviral vector. LICh pseudotyped reporter viruses were generated in 293T cells co-transfected with LICh dual reporter genome vector, SIV3+ packaging vector, JRFL envelope encoding plasmid and REV expression plasmid as described in Methods.

Two sets of depo provera-treated PTMs were vaginally inoculated with the LICh vector (10^5^-10^6^ TCID50) 28 days (pilot study) or 25 days (preclinical study) post TDF-IVR insertion (Figure 1). The first set of animals was used for pilot analysis (pilot study) and included 4 TDF-IVR-treated PTMs (BB432, BB588, BB401 and BB925) and 2 control PTMs with no IVR (BB966 and 96P047). Animals were euthanized 48 hrs after vector challenge. The second set of animals were used to perform in depth PD/PK (preclinical study) and included 6 TDF-IVR-treated PTMs (BB548, 1.6348, 1.8678, BB187, BB963, BB535) and 1 no-IVR control (BB529). All the animals in the second group were euthanized 72 hours post viral inoculation to potentially increase the detectable luciferase signal (Figure 1). The entire intact FRT was collected at necropsy and examined for luciferase activity using an In Vivo Imaging System (IVIS). We first identify and define unspecific background signal by IVIS evaluation in the absence of exogenous luciferin. The tissue was soaked in d-Luciferin, while ovaries were injected with d-Luciferin, and reimaged to identify specific luciferase activity. Positive signal was found in foci throughout the entire FRT including vagina, cervix, uterus, ovaries, and inguinal lymph nodes in both 48 hours control animals (96P047 and BB966) from the pilot study. However, no signal was detected in the control animal (BB529) of the preclinical study necropsied 72hrs after vector challenge.

**Figure 1.**
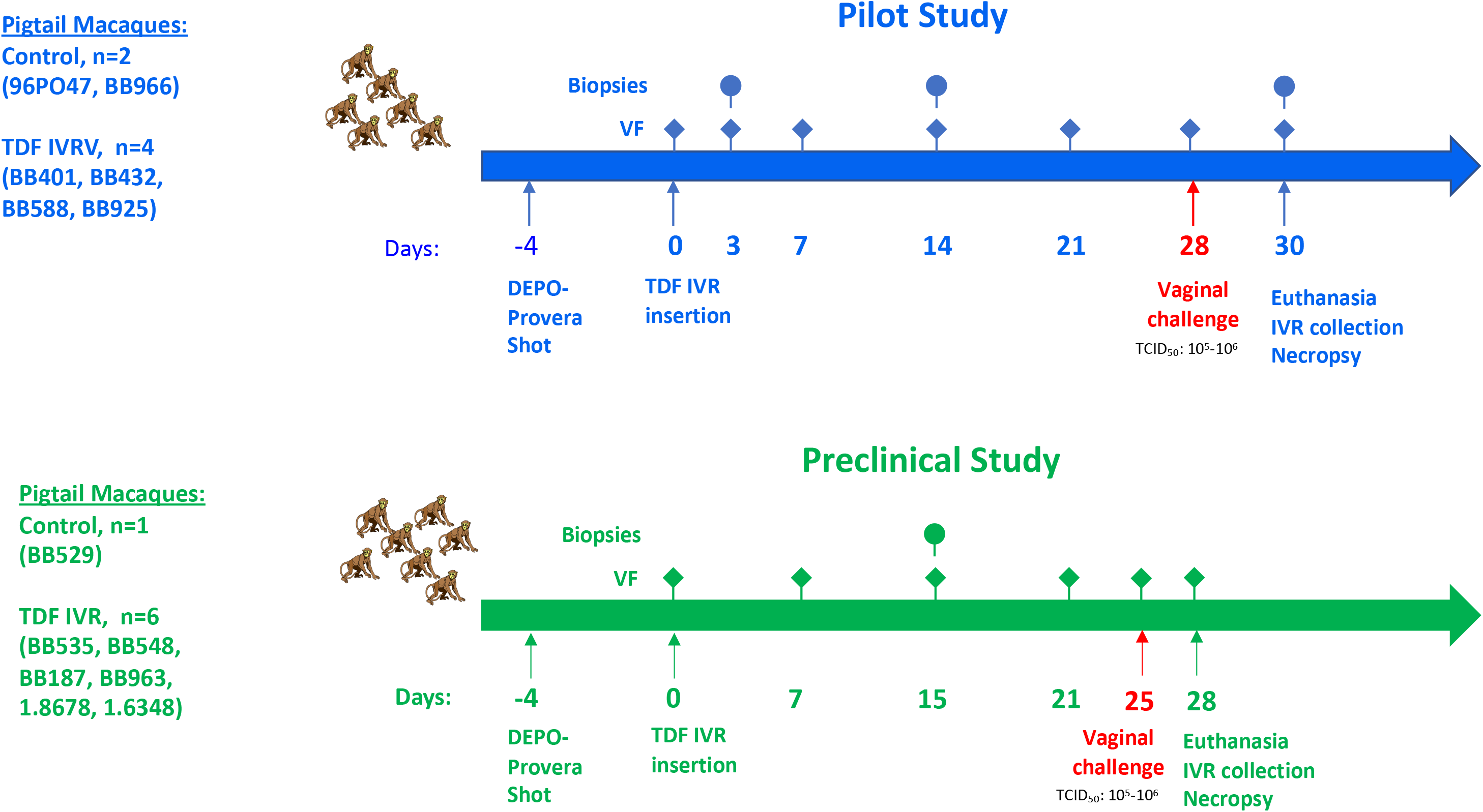
Timeline of the studies. All animals were treated with DEPO Provera contraceptive 4 days prior to the TDF-IVR insertion. Animals in the pilot study (96P047, BB966, BB401, BB925, BB432, BB588) were challenged with SHIV reporter virus 28 days after ring insertion and euthanized on day 30. Animals in the preclinical study (BB529, BB535, BB548, BB187, 1.8678, 1.6348, BB963) were challenged with SHIV reporter virus 25 days after ring insertion and euthanized on day 28. FRT processing occurred one day after necropsy for both studies.

Luciferase activity in the control animal 96P047 is shown in Figure 2A. The FRT was then dissected to separate different FRT regions and further cut into smaller pieces that were reimaged for luciferase activity and then embedded into Optimal Cutting Temperature (OCT) media. Luciferase signal persisted in several dissected tissue pieces as shown in Figure 2B. This result is in accordance with our previous study showing the susceptibility of the entire FRT to viral entry (17). In the animals that were treated with the TDF-IVR, we detected very low or no luciferase activity in the lower FRT in both the pilot and preclinical studies. However, luciferase signal was present in the ovaries of six out of the ten TDF-IVR- protected animals (Table 1). Moreover, signal was also evident in the lymph nodes of three of the ten TDF-IVR-treated animals (Table 1). Luminescence in the FRT of a representative animal BB548 is shown in Figure 2C-D. The FRT tissue of all TDF-IVR-treated animals was further dissected into smaller pieces and reimaged as described above for controls. No additional luminescence was identified in smaller tissue pieces as shown in Figure 2D for animal BB548. All tissues were stored in OCT media at -80°C for further analyses. Luciferase activity throughout the FRTs and lymph nodes of all animals is summarized in Table 1.

**Figure 2.**
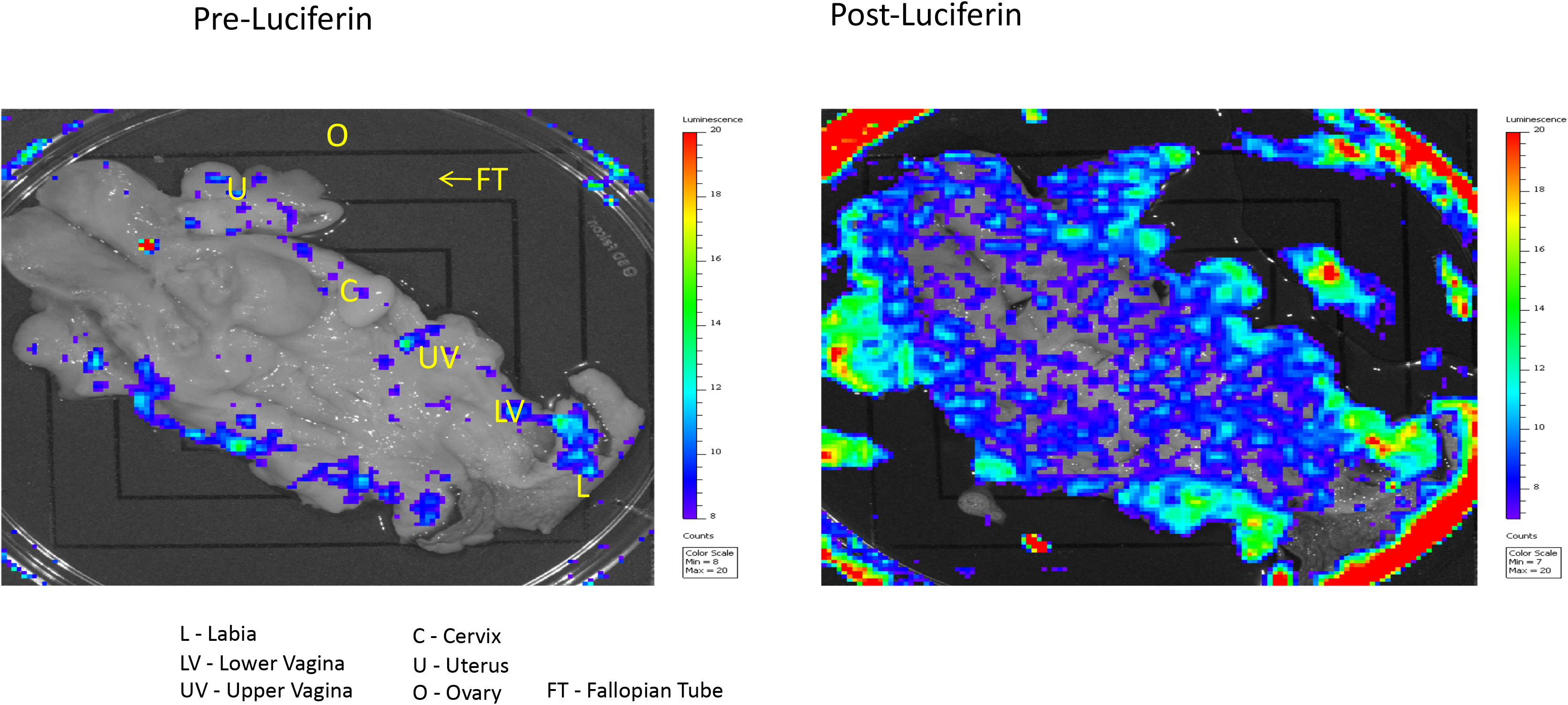
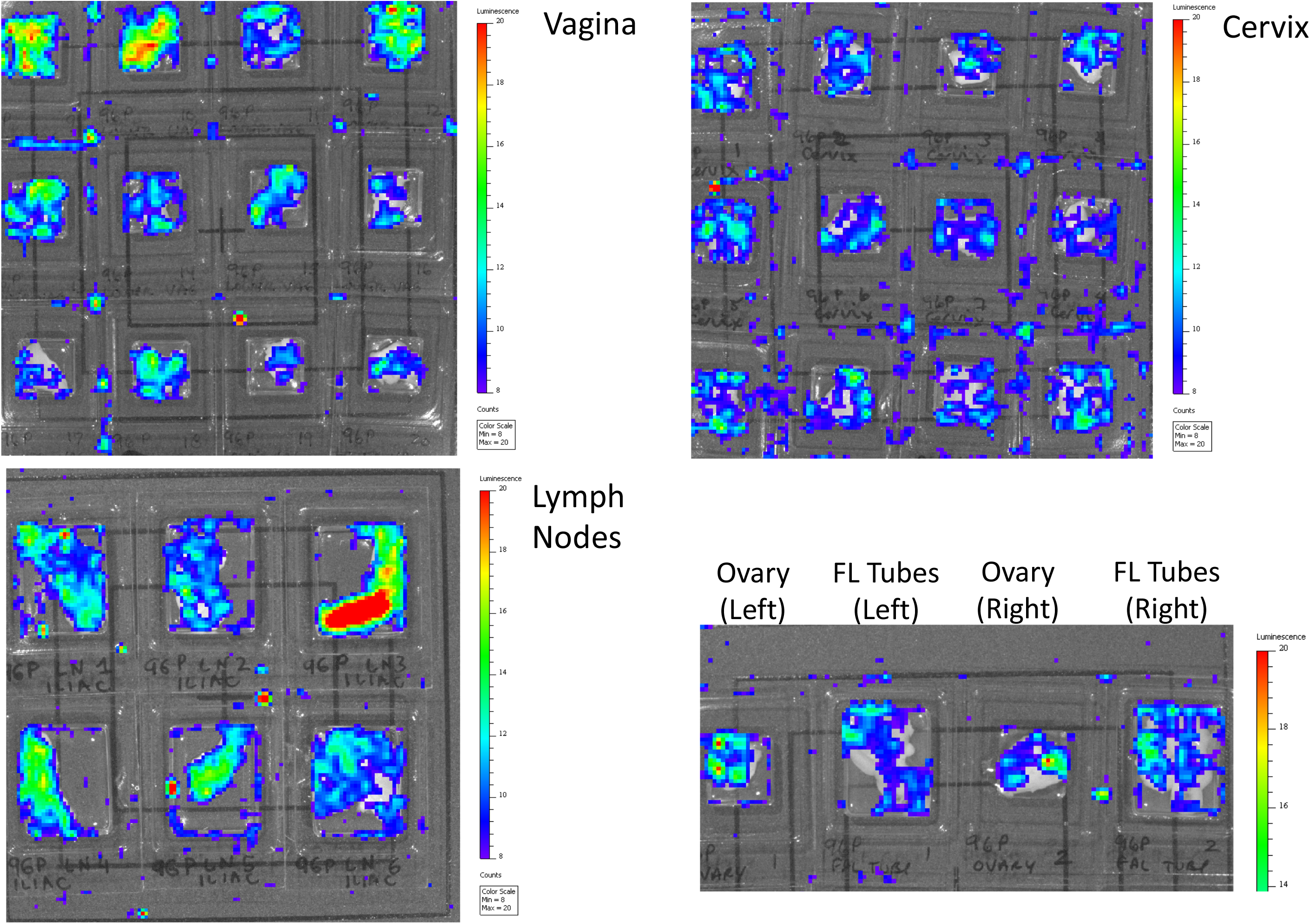
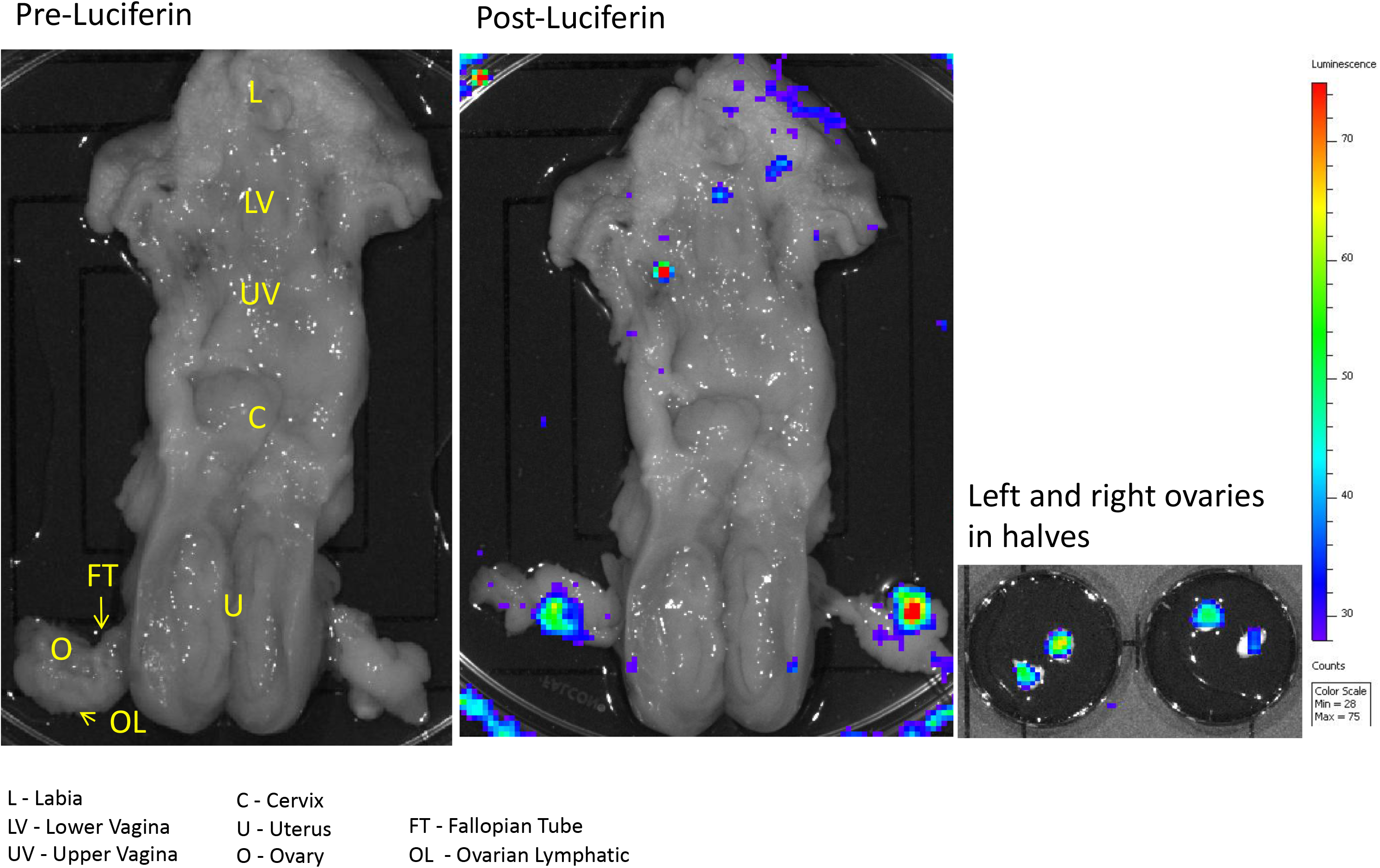
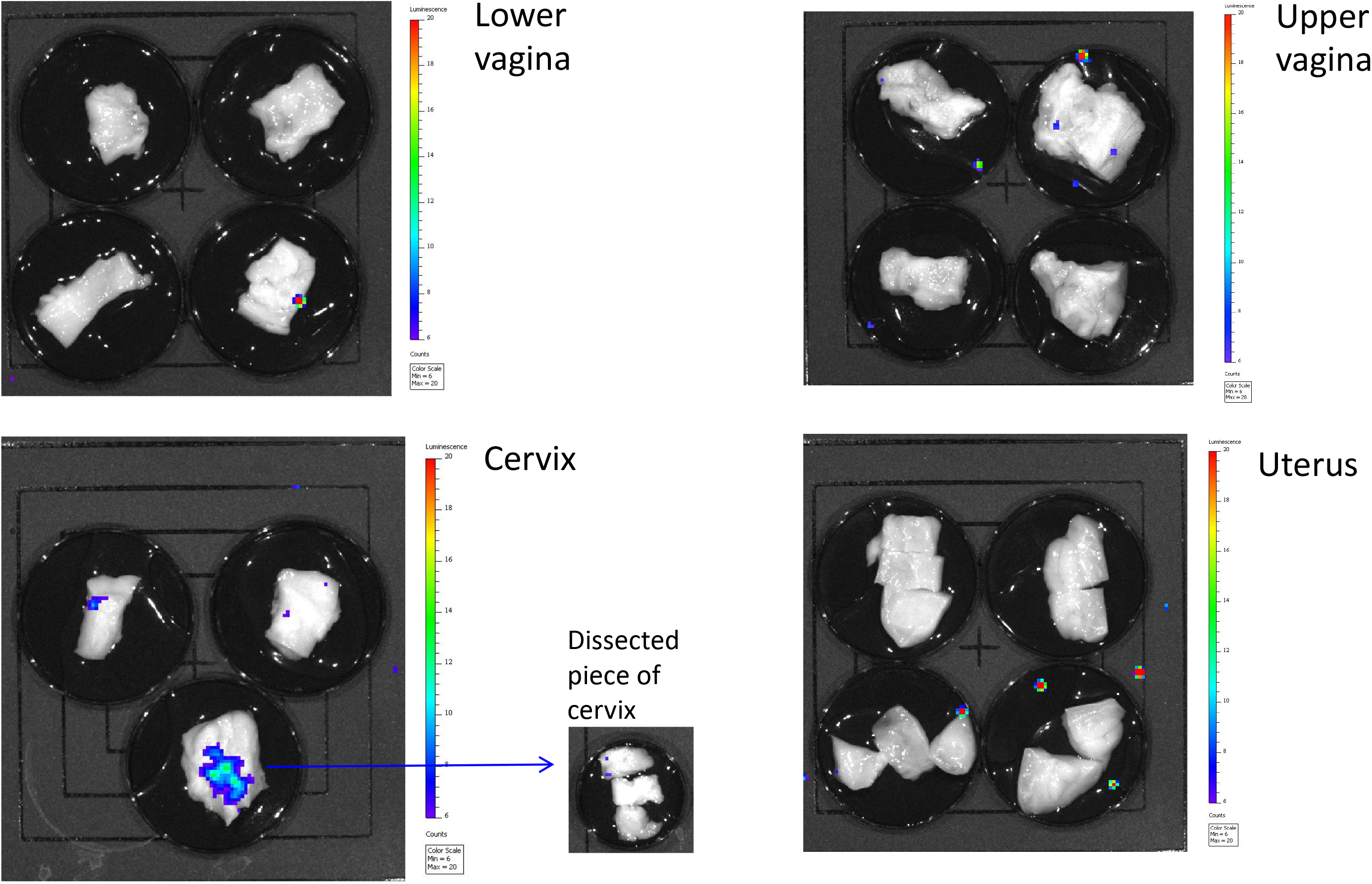
Luciferase activity in female reproductive tract of pigtail macaques challenged with a single high-dose reporter virus expressing Luciferase and mCherry genes. Luciferase activity was induced by soaking the tissue in luciferin and detected using IVIS. **(2A).** Positive signal was identified throughout the FRT in the control animal 96P047 (Post-luciferin image). **(2B).** Tissue of the 96P047 animal was dissected into smaller pieces and reimaged. **(3C).** Whole FRT scan of TDV-IVR treated animal BB548. Positive signal was identified in both ovaries. **(2D)**. Dissected and reimaged tissue of TDV-IVR treated animal BB548.

**Table 1.**
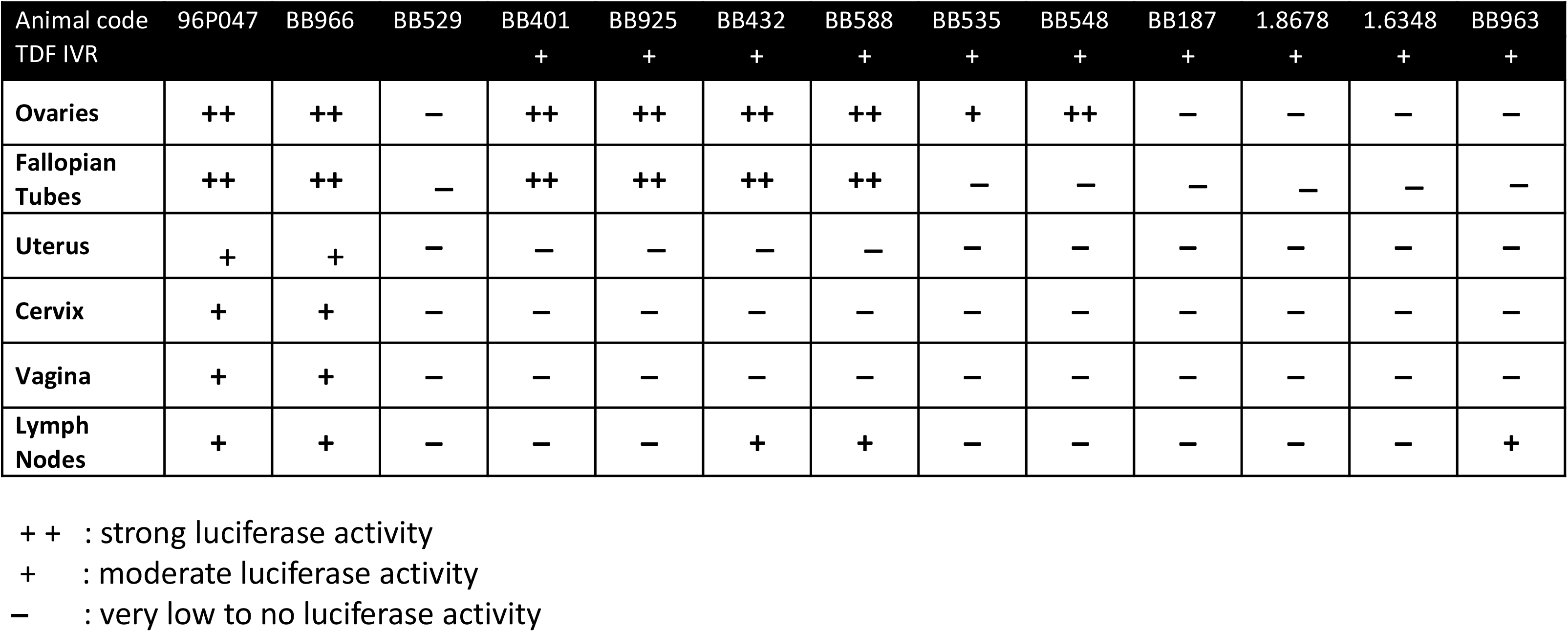
Summary of luciferase reporter expression throughout the FRT and lymph nodes of pigtail macaques analyzed.

### Single round infectious virus transduced CD4+ immune cells in ovaries and lymph nodes

An ovary of the control animal BB966 (Figure 3A) and iliac lymph node of the control animal 96P047 (Figure 3B) were analyzed for the presence of transduced cells by fluorescent microscopy (Figure 3A, B). mCherry and luciferase positive cells were identified in both tissues in accordance with the strong luciferase activity observed using IVIS. Since the ovaries of TDF-IVR-treated animals also presented strong luciferase activity with IVIS, we analyzed the ovaries of the animals in the preclinical study further by fluorescent microscopy (Figure 3C, D). We observed multiple areas with a high auto fluorescence background that is common in ovaries. Thus, we validated the signal using spectral imaging (maximum emission at around 610 nm for mCherry and 665 nm for luciferase in CY5, Supplemental Figure 3A, B) and used this criteria to distinguish between non-specific and specific signal to identify cells transduced by the SHIV-LICh pseudovirus.

**Figure 3.**
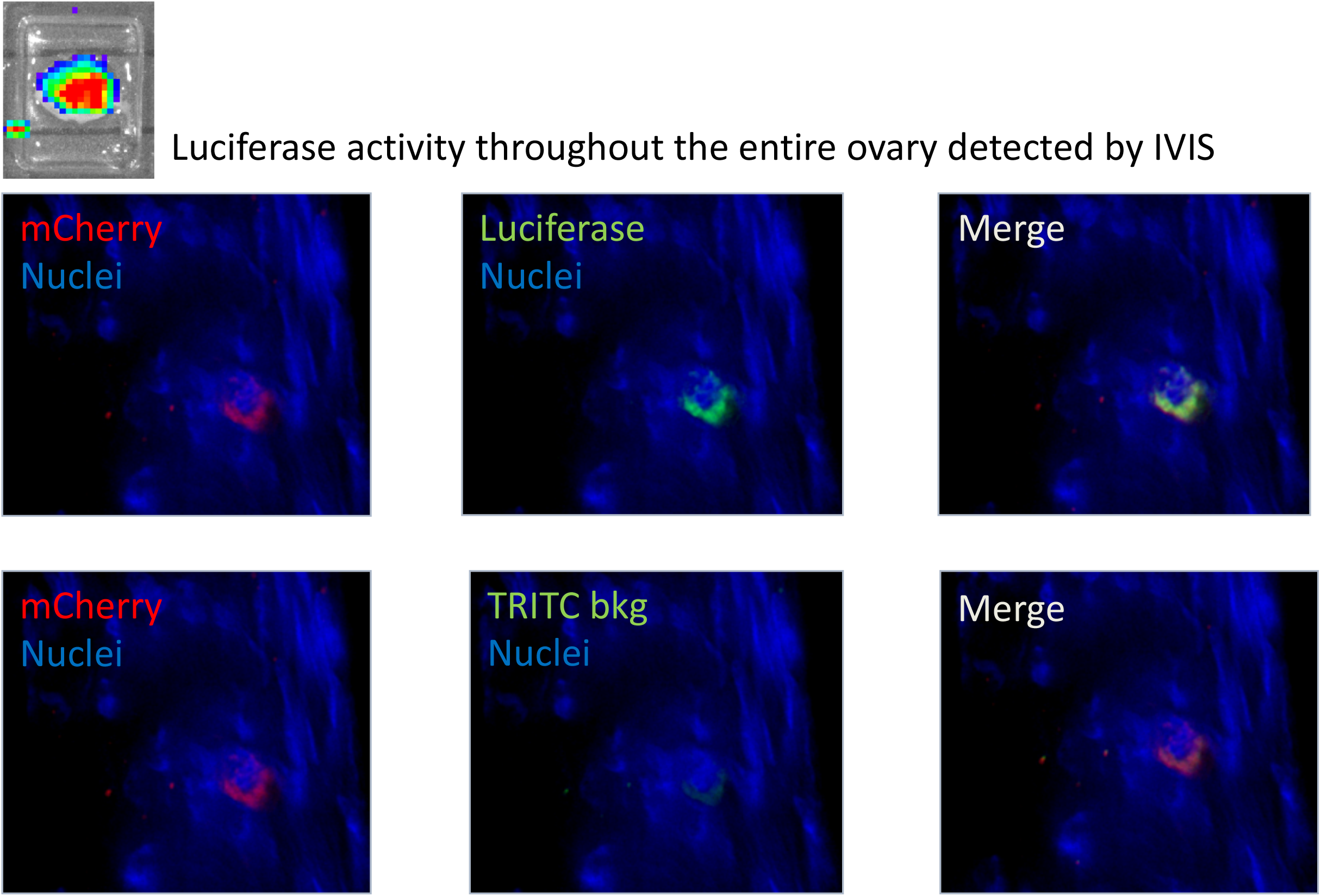
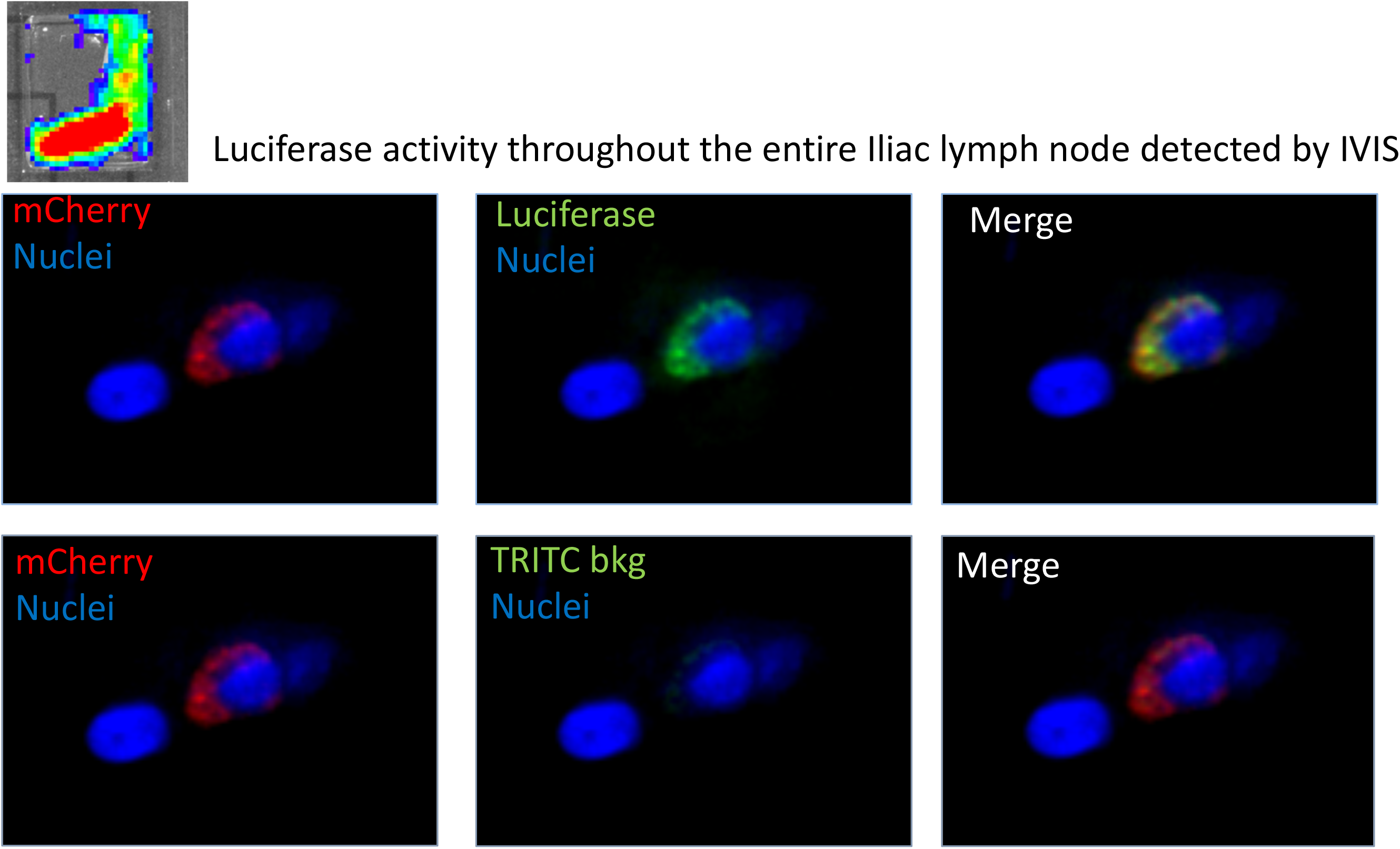
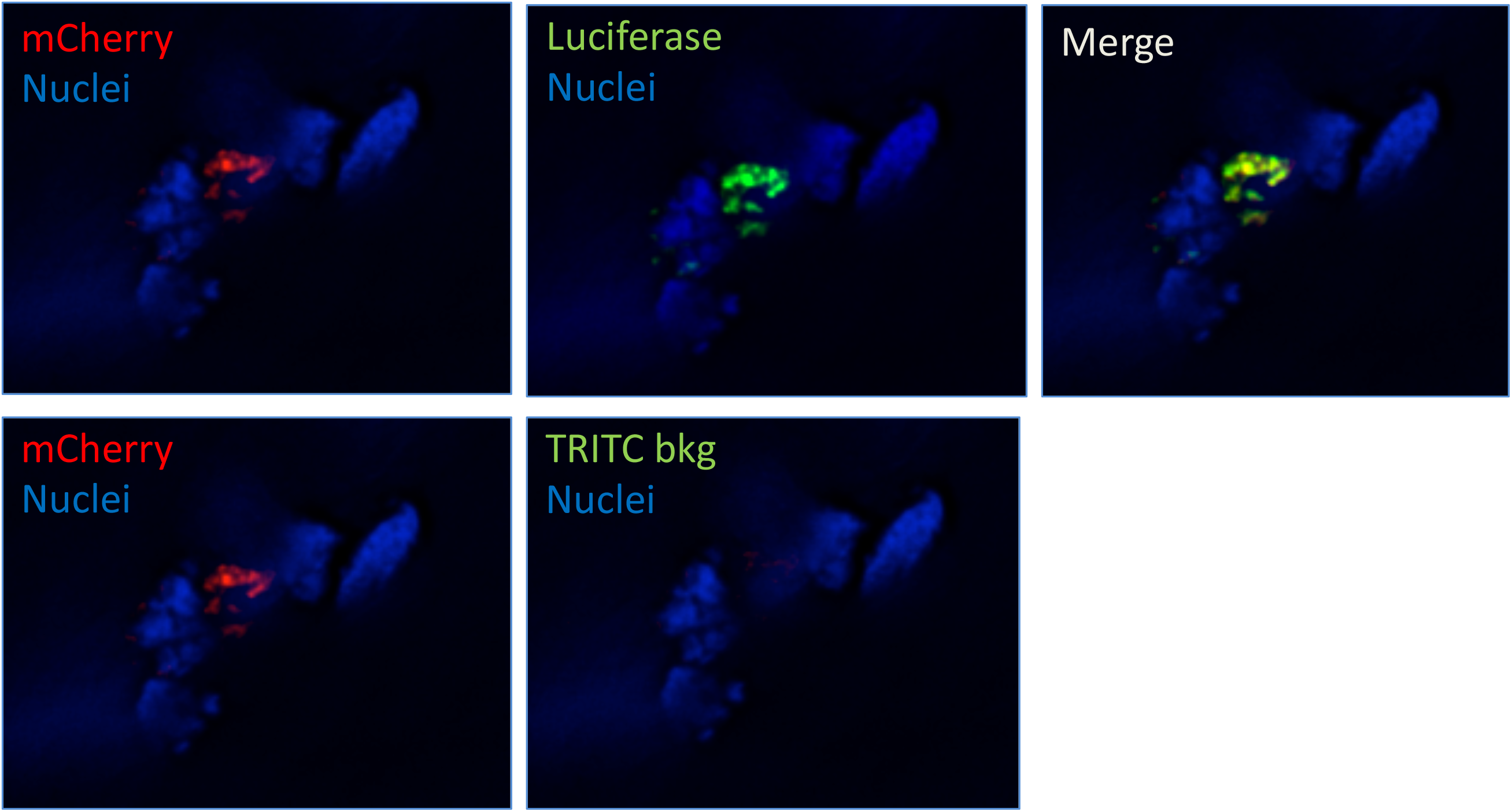
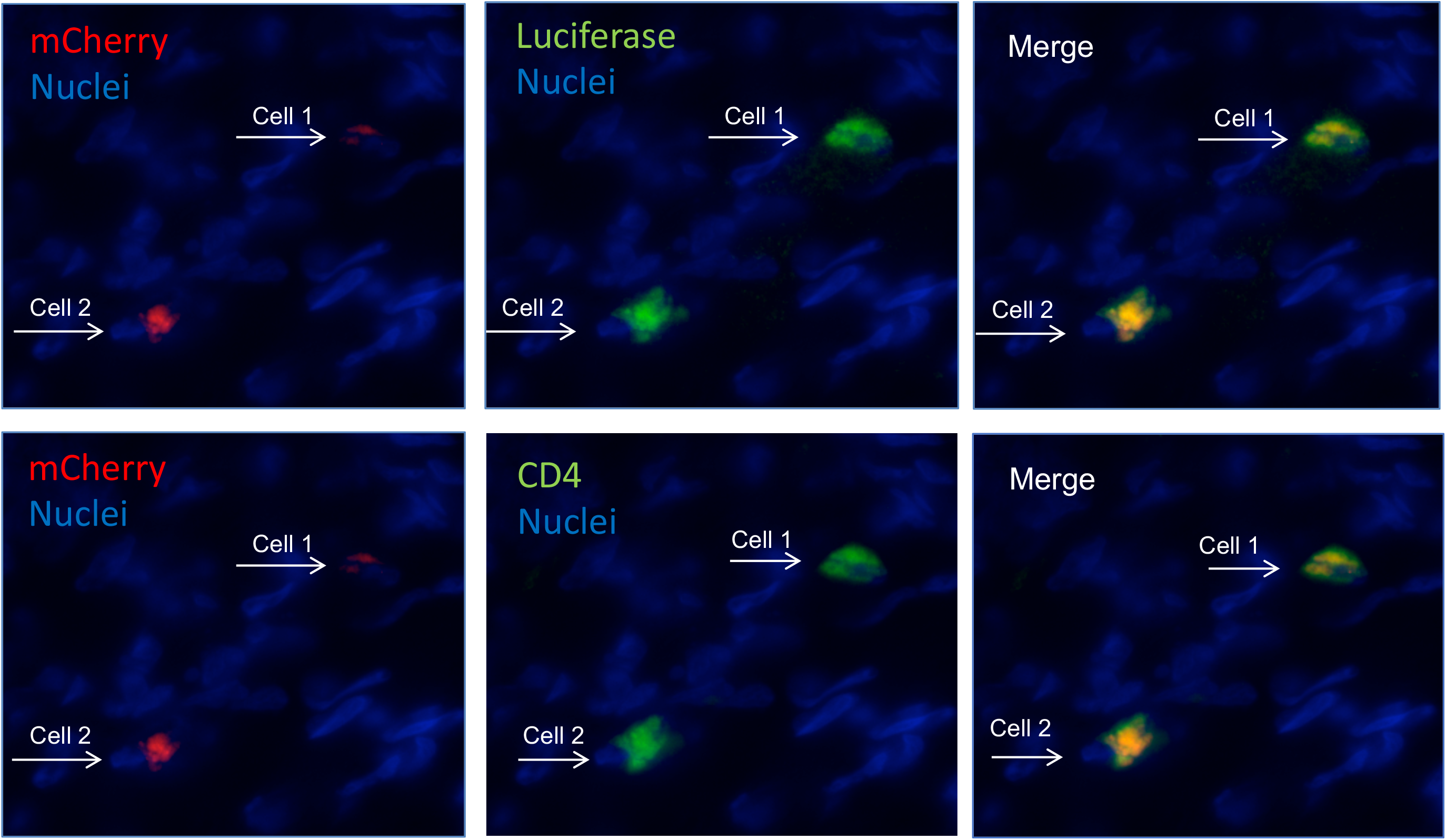
mCherry and Luciferase expression in the tissue of female reproductive tract of challenged pigtail macaques. IVIS re-imaged tissue pieces were cryosectioned and immunostained with anti-Luciferase antibody and DAPI. Immunostained tissue sections were surveyed using fluorescence microscopy. Luciferase (green) and autofluorescent mCherry (red) positive cells were identified in right ovary of the BB966 control animal (**3A)**, Iliac lymph node of the 96P047 control animal (**3B),** ovarian tissue of TDF-IVR treated animal BB535 (**3C**), and ovarian tissue of TDF-IVR treated animal BB548, where transduced cells were phenotyped with anti-CD4 antibody (**3D**). DAPI was used to stain nuclei (blue). Specificity of transduction was confirmed by low fluorescent intensity in TRITC channel.

The transduced cells were phenotyped by co-staining the tissue with anti- CD4+ antibody. This analysis demonstrated that some cells double positive for mCherry and luciferase also expressed the CD4+ receptors (Figure 3D). Frequently, more than one transduced cell was found at locations that were rich in CD4+ cells. Such an example is presented in Figure 3D. We identified a median of 2 (range 0-8) of transduced cells in the ovaries of the 6 TDF-IVR treated animals and 2 cells identified in the control animal (Table 2). The 1.8678 animal was an exception; no mCherry/luciferase positive cells were present in cryosections of both ovaries, while transduced cells were identified in both ovaries of the animal number BB535.

**Table 2.** Number of transduced cells (mCherry/luciferase positive) identified in ovaries of SHIV challenged pigtail macaques that were administered with the TDF-IVR.

| Animal<br>code:<br>TDF IVR | BB529 | BB535 | BB548 | BB187 | 1.8678 | 1.6348 | BB963 |
| --- | --- | --- | --- | --- | --- | --- | --- |
|  | - | + | + | + | + | + | + |
| Ovary 1 | 2 | 8 | 2 | 1 | 0 | 4 | 1 |
| Ovary 2 | - | 2 | 0 | 0 | 0 | - | - |

Number of cells found per ovary
- : Tissue not available for screening

### Pharmacokinetics of the TDF-IVR

Pigtailed macaques were used for both the pilot study and the preclinical study as shown in Fig 1. The TDF-IVR used was the same that resulted in full protection in a previous challenge study (13). To ensure we were getting TFV levels similar to the efficacy study, TFV was measured in vaginal tissue biopsies both proximal and distal to IVR placement, and in vaginal fluids (VF). The TFV tissue levels were measured from biopsies collected on days 3, 14 and 30 for the pilot study and from day 15 biopsies in the preclinical study (Figure 4A). When the IVR was present the TFV levels were around 10^5^ ng/g of tissue proximal to the IVR placement whereas the distal tissues were about a log lower. Two days after removal of the IVR the tissue levels dropped 1-2 logs below from what was observed on day 14/15 of the study but were still approximately 10^3^ ng/g of tissue. TFV-diphosphate (TFV-dp), the active intracellular species of TFV, was measured in whole biopsies for the preclinical group on day 15, and from lymphocytes isolated from vaginal tissue two days after ring removal in the pilot study (Figure 4B). These values were quite variable (range 10-1000 fmol/mg of tissue and 80-800 fmol/10^6^ isolated lymphocytes) but provide evidence that the TDF is readily converted to TFV-dp in vaginal tissues. The presence of TFV-dp two days after IVR removal suggests that some of these cells are retained in the vaginal tissue and are not trafficking to other tissues. The median TFV levels in VF proximal to the IVR were consistently around 10^6^ ng/mL whereas the median TFV in samples collected distal to the IVR ranged from 10^4^ – 10^6^ (Figure 4C). These levels are consistent with those reported for the efficacy study (13).

**Figure 4.**
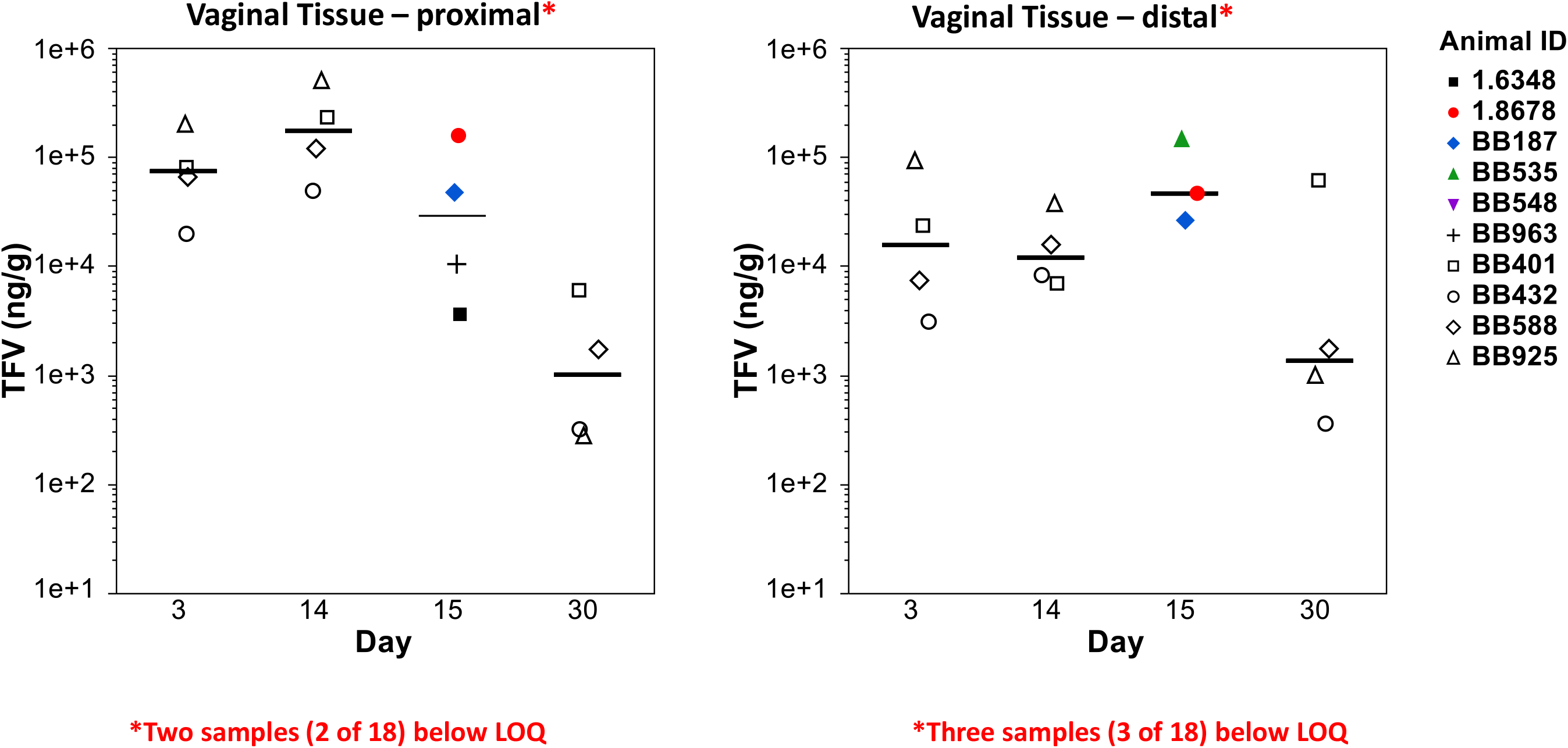
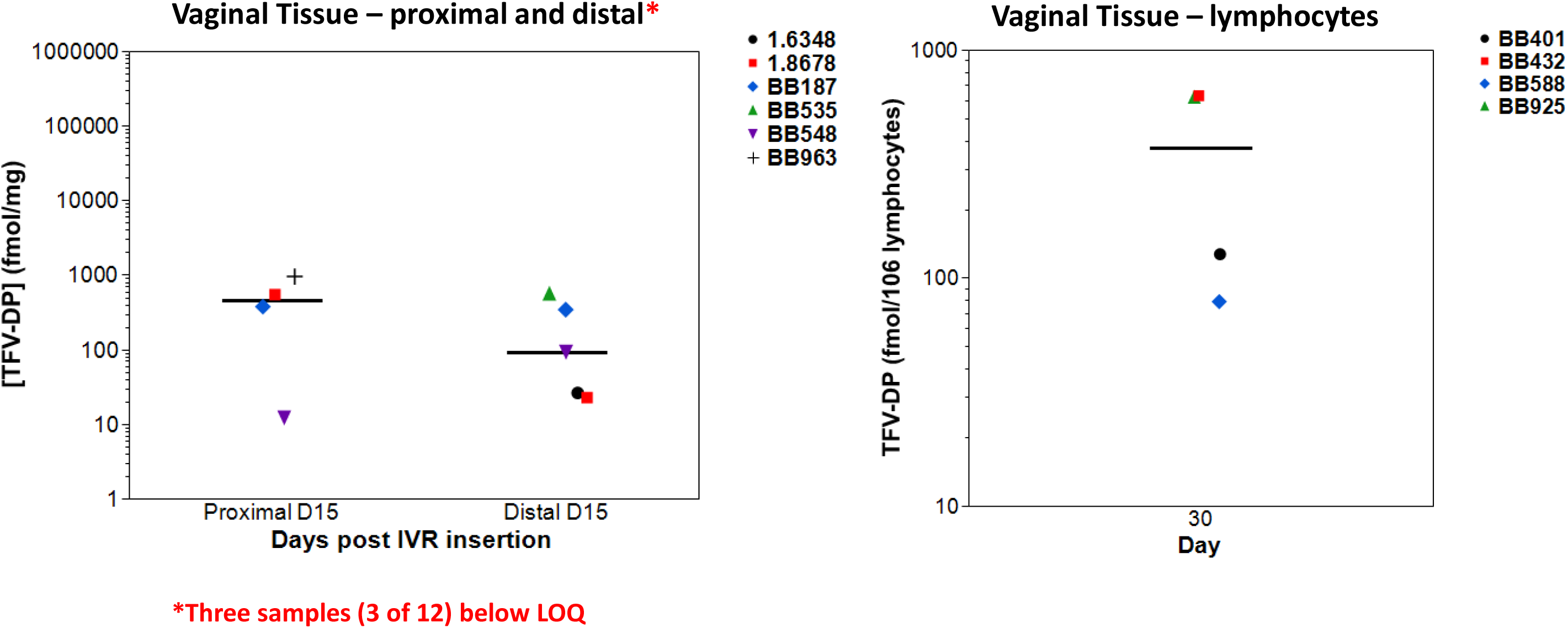
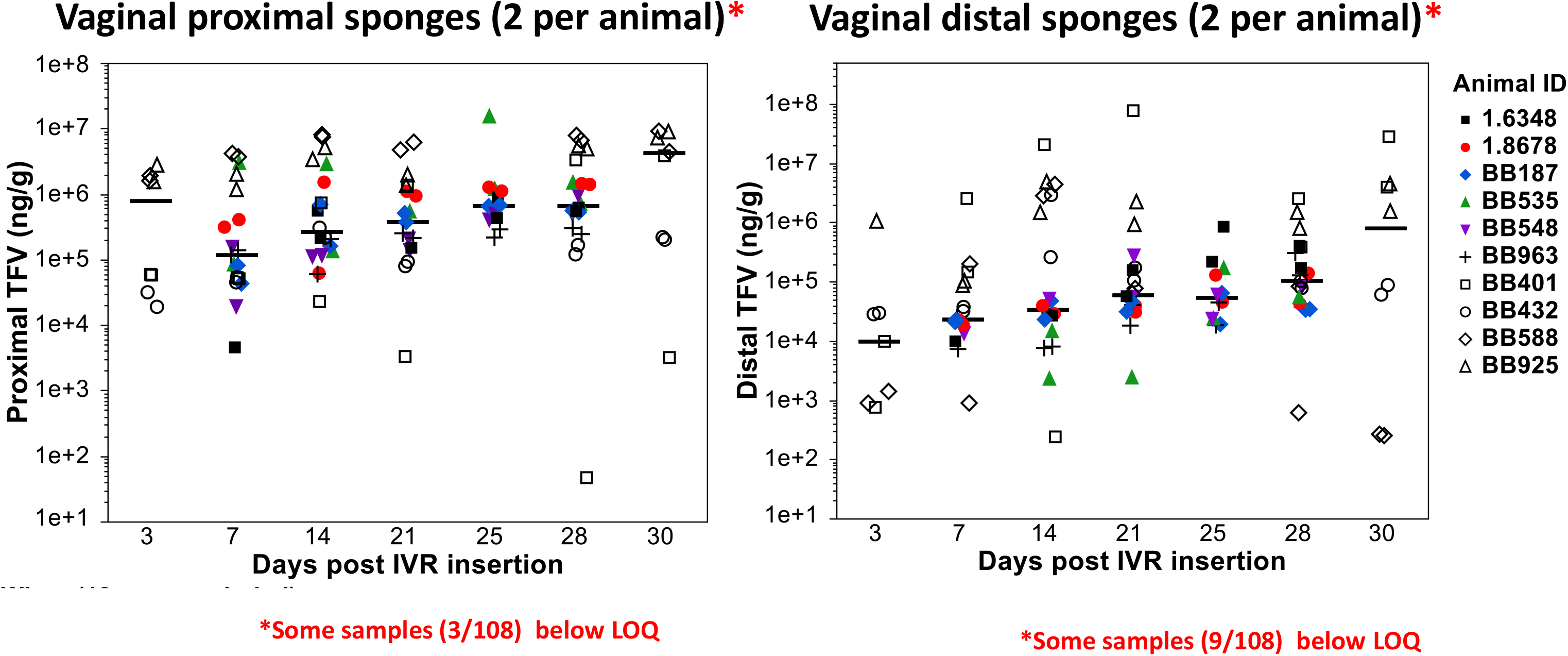
**Drug levels during treatment with IVR 4A**. TFV levels as measured by LC-MS/MS from vaginal tissue biopsies collected on days 3 and 14 post IVR insertion, and vaginal tissue sections collected at necropsy (day 30) for the pilot study. In the preclinical study biopsies were collected on day 15. **4B**. TFV-DP concentrations were determined in whole tissue (day15) and leukocytes isolated from processed vaginal tissues (day30). **4C**. TFV levels from sites proximal and distal to IVR placement.

TFV levels at necropsy for the animals in the preclinical study were measured throughout the FRT and ranged from below LLOQ to 5000 ng/g tissue with a strong dependence on the proximity of the tissue to the placement of the IVR (Figure 5). Statistically significant location dependence of tissue drug levels was observed using the Wilcoxon rank sum test (P < 0.0001). Further pairwise testing using the Wilcoxon method demonstrated that the higher drug levels in the cervix and upper vagina were not different (P = 0.39), but that drug levels in the upper vagina and lower vagina were different (P = 0.01), as were the drug levels in the upper reproductive tract (uterus, ovaries, fallopian tubes) compared to the upper vagina/cervix (P = 0.01) (Figure 5). Thus, our results demonstrate the gradient of tissue drug levels emanating from the site of the ring (Figure 5 & 7). We observe lower drug levels in the upper reproductive tract for all animals. Drug levels in the ovaries, where transduced cells were found in five of the six IVR-treated animals, ranged from below our limit of quantitation (50 ng TFV/g tissue) to 400 ng TFV/g tissue (Figure 5). The measured levels of TFV throughout the FRT of the animals in the preclinical study at necropsy were consistent with the levels of TFV in the vaginal tissues of the macaques from the pilot study collected 48-hrs after SHIV-LICh challenge (Figure 4A).

**Figure 5.**
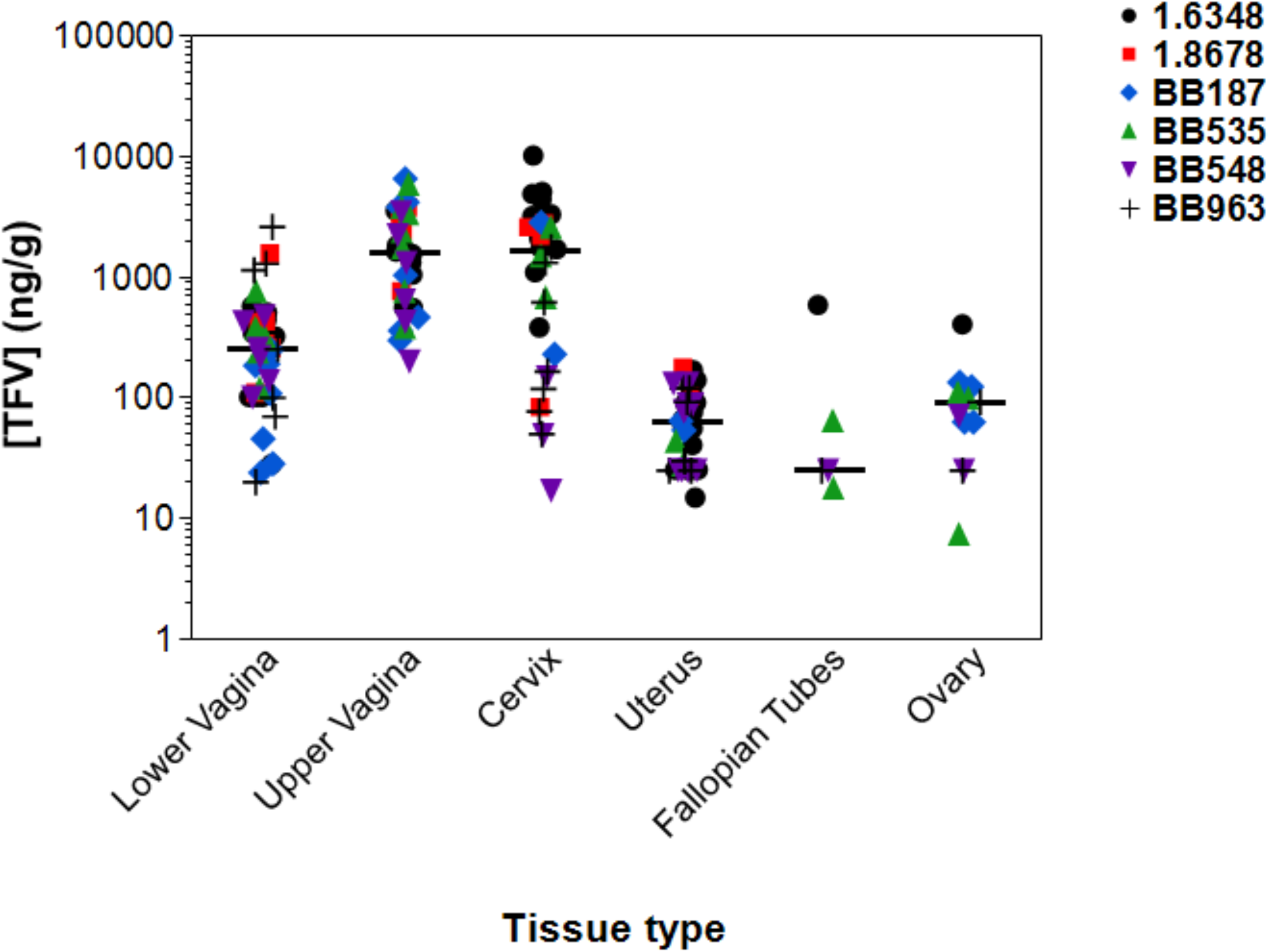
Post-necropsy TFV levels in macaque tissue. TFV levels as measured by LC-MS/MS at necropsy for the preclinical study as a function of location in the FRT.

### Complete screen of FRT using nested PCR reveals uneven inhibition of viral reverse transcription by TDF-IVR

We utilized nested PCR methods to survey genital tissues of all animals in the pre-clinical study (necropsied 72-hours after SHIV-LICh challenge) for proviral DNA detection. Because animals had been infected with the SHIV162p3 prior to the high- challenge with LICh vector we designed PCR primers to target IRES and WPRE DNA elements and mCherry gene, all specific for our viral reporter genome to avoid cross-detection of SHIV163p2 LTR sites in these animals. Outer PCR products were amplified between IRES and WPRE elements, while inner PCR products were generated from the mCherry gene located between IRES and WPRE. Scheme of primer design is represented in Supplemental Figure 1.

To evaluate nested PCR sensitivity, we infected 293T cells with SHIV-based virus containing the dual reporter genome. As described in methods, infected cells were selected based on expression of mCherry protein using flow cytometry. Then, mCherry expressing 293T cells were lysed for DNA isolation. DNA was separately extracted from non-infected 293T cells. Genomic DNA of both cell types, mCherry positive and 293T wild type cells, were mixed together in a ratio that corresponds to 100 mCherry positive cells in 250 ng of total DNA per PCR reaction which represents approximately 3.8 x 10^4^ cells. Serial dilutions of mixed genomic DNAs were prepared using genomic DNA of wild type cells to test efficiency of the primers. The lowest DNA dilution corresponded to less than 1 infected cell per PCR reaction. In this primer efficiency assay we demonstrated high sensitivity of this method and were able to detect as low as 3 copies of proviral DNA per 250 ng of genomic DNA (Supplemental Figure 2). As a positive control we separately amplified LTR elements and we only used these sites for the PCR sensitivity assay, however not for the screening of animal tissue. The dual reporter genome contains two LTR elements (Supplemental Figure 1), and we therefore observed higher PCR sensitivity resulting in detection of less than one copy of proviral DNA in a single PCR reaction of 293T genomic DNA (Supplemental Figure 2).

Our PCR data show that early viral events occurred in an ovary and vaginal and cervical tissue of the control animal (BB529) (Figure 6A, Table 3) despite detecting no luciferase activity in these tissues (Table 1). This result suggested that we could find rare viral events by using a highly sensitive nested PCR method. In support of this, viral transduction was detected by PCR in an ovary of the IVR treated animal (BB187) that did not demonstrate luciferase activity (Figure 6A, Table 1). Overall, we detected mCherry DNA in the ovaries, ovarian lymphatic, fallopian tubes, cervix, vagina and lymph nodes of studied animals. These data are summarized in the Table 3 and Figure 6B. Five animals (BB963, BB548, BB187, 1.6348 and 1.8678) of six TDF-IVR treated were positive for the presence of proviral DNA in cervical and vaginal tissue although none of them was positive for luciferase activity (Table 1). Proviral DNA was also detected in the ovaries of two animals (BB187 and 1.8678) (Table 3) that were negative for luciferase signal (Table 1). On the other hand, no viral transduction was identified in ovaries of two animals (BB548 and BB535) (Table 3) that were positive in the assay of luciferase activity (Table 1). Since only a few sections of ovaries were used for nested PCR evaluation it is very likely that pseudoviral transduction did not occur in these sections. Ovarian lymphatic of two animals (BB548 and 1.8678) and the fallopian tube of an animal (BB548) were also positive for proviral DNA (Table 3). These data suggest that TDF-IVR may not completely protect the genital tract of pigtail macaques from a single high viral dose challenge and demonstrate the location of the initial sites of viral transduction. The nested PCR data suggest that the highest frequency of proviral DNA appears in cervical and vaginal tissue sections (Table 3) of TDF-IVR animals.

**Figure 6.**
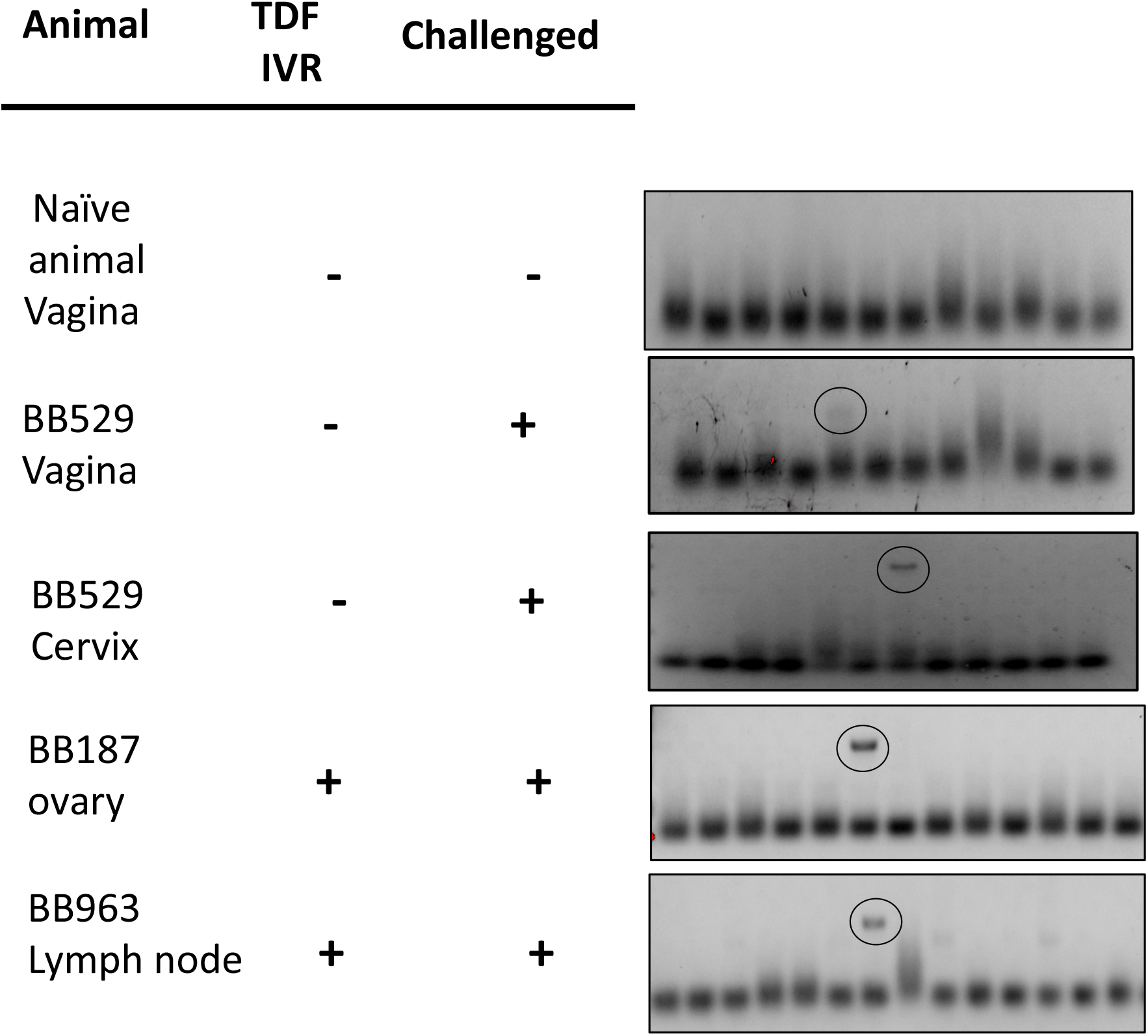
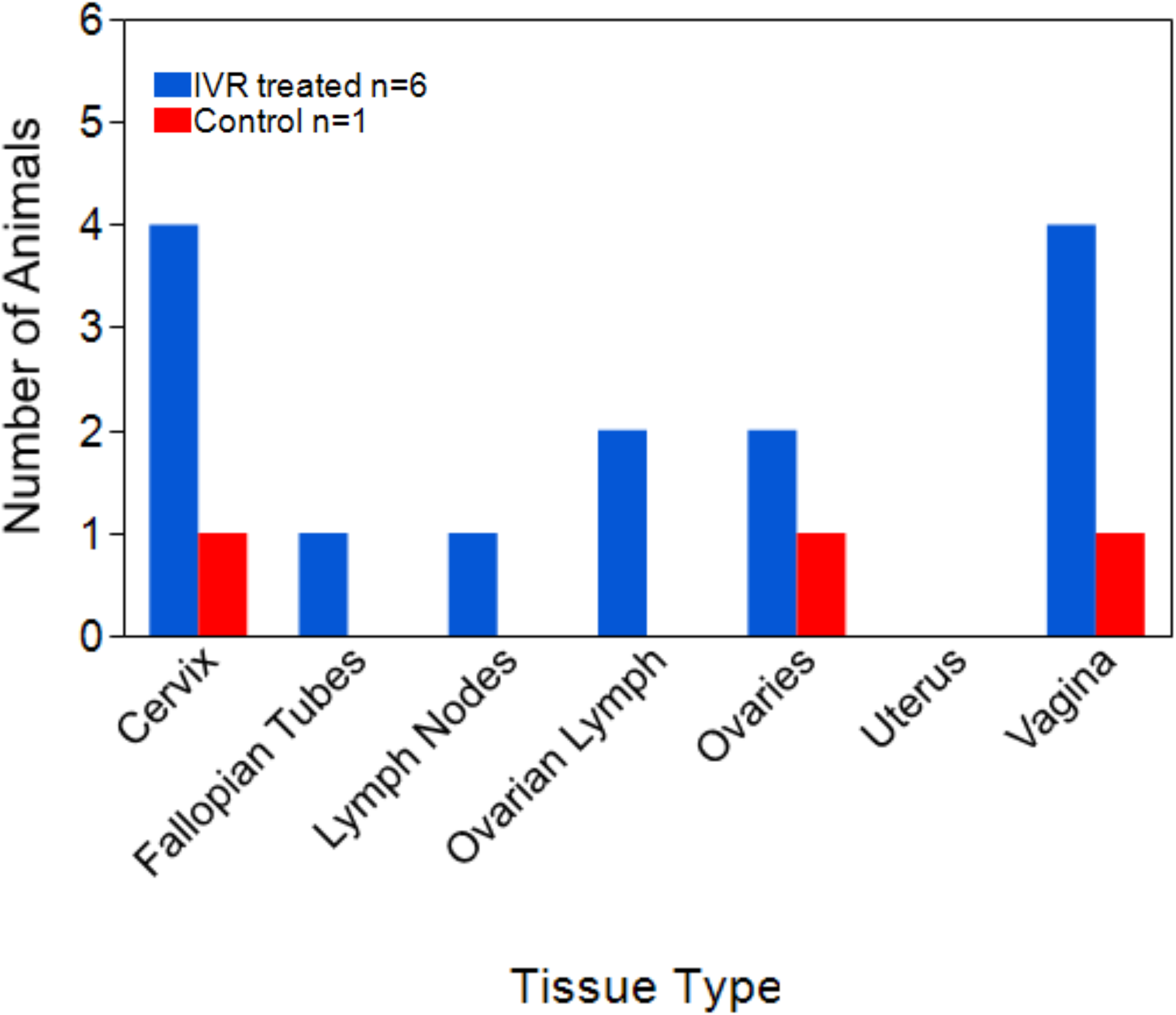
Nested PCR screening results. Dissected tissue of the entire FRT and lymph nodes were cryosectioned for DNA extraction following nested PCR. Each dissected tissue piece was surveyed for the presence of proviral DNA (mCherry gene) copy in total of 250 ng DNA per reaction in at least 12 PCR reactions. **(6A**) 250 bp long PCR products visualized in gel and sequenced using mCherry primers for naïve animal, control animal BB529 (vagina and cervix), and IVR-treated animals BB187 (ovary) and BB963 (lymph node). (**6B**) Summary of proviral DNA detection using nested PCR throughout FRT and lymph nodes.

**Table 3.**
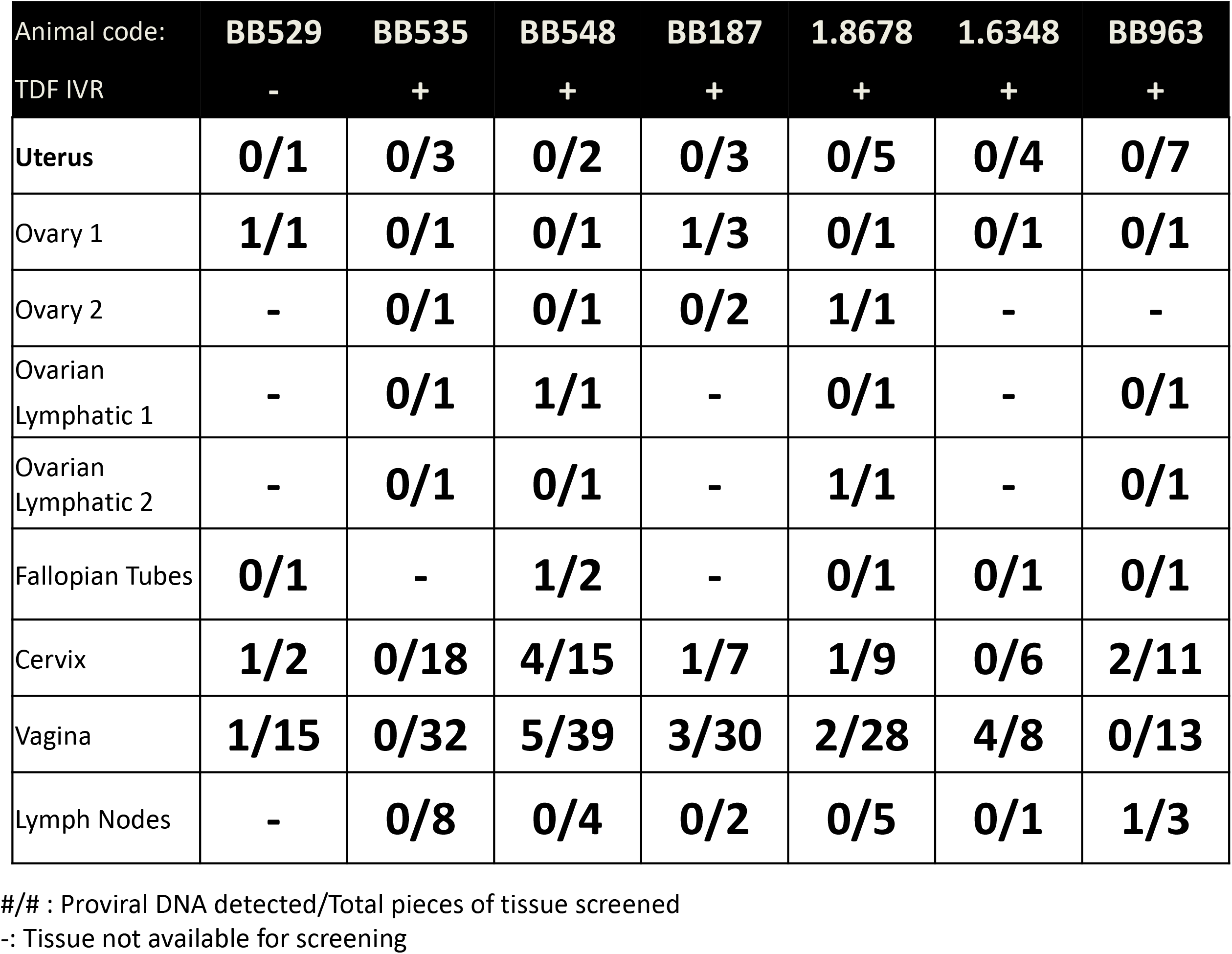
Number of proviral DNA detected throughout the entire FRT and lymph nodes of Pigtail Macaques challenged with a single high dose of SHIV.

| Animal code: | **BB529** | **BB535** | **BB548** | **BB187** | **1.8678** | **1.6348** | **BB963** |
| --- | --- | --- | --- | --- | --- | --- | --- |
| TDF IVR | - | + | + | + | + | + | + |
| Uterus | **0/1** | **0/3** | **0/2** | **0/3** | **0/5** | **0/4** | **0/7** |
| Ovary 1 | **1/1** | **0/1** | **0/1** | **1/3** | **0/1** | **0/1** | **0/1** |
| Ovary 2 | - | **0/1** | **0/1** | **0/2** | **1/1** | - | - |
| Ovarian Lymphatic 1 | - | **0/1** | **1/1** | - | **0/1** | - | **0/1** |
| Ovarian Lymphatic 2 | - | **0/1** | **0/1** | - | **1/1** | - | **0/1** |
| Fallopian Tubes | **0/1** | - | **1/2** | - | **0/1** | **0/1** | **0/1** |
| Cervix | **1/2** | **0/18** | **4/15** | **1/7** | **1/9** | **0/6** | **2/11** |
| Vagina | **1/15** | **0/32** | **5/39** | **3/30** | **2/28** | **4/8** | **0/13** |
| Lymph Nodes | - | **0/8** | **0/4** | **0/2** | **0/5** | **0/1** | **1/3** |

#/# : Proviral DNA detected/Total pieces of tissue screened

The PCR-data from the 6 PTMs treated with the TFD-IVR in the preclinical study were overlayed to the IVIS images of the whole FRTs to present early viral events in a form of 2D FRT maps (Figure 7). To this, the TFV levels obtained from each corresponding piece of tissue were overlayed to the PCR data to visualize the activity of the TDF-IVR at the anatomical level. These FRT maps show the highest drug levels in the middle of the FRT, where the ring was located (upper vagina, endocervix) across all six preclinical animals. Drug levels decreased in upper (uterus, fallopian tubes, ovaries) and lower (labia, lower vagina) FRT regions that were distal from the TDF-IVR location. Furthermore, plus/minus signs on the maps describe positive/negative nested PCR detections indicating random viral transductions throughout the FRTs regardless of drug levels detected. The number of detected transduced cells in ovaries are noted over each animal’s ovary.

**Figure 7.**
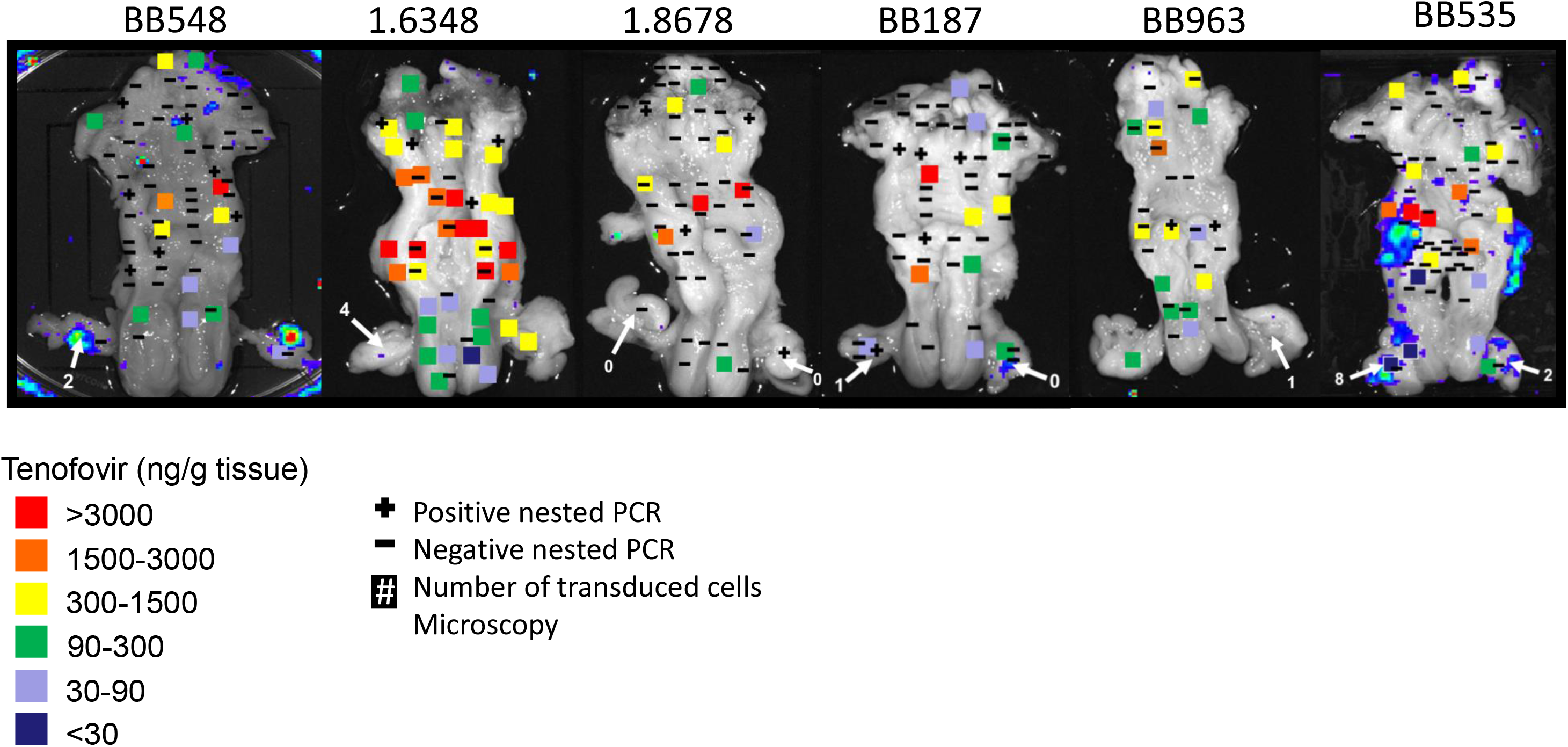
Summary of FRT post-necropsy analyzes. FRT 2D-maps of animals administered with TDF-IVR in preclinical study. Pigtail macaques were treated with TDF- IVR for 28 days. On day 25 post TDF-IVR insertion animals were vaginally challenged with a single high viral dose (10^5^-10^6^). 72 hrs later animals were scarified and FRTs were analyzed using three different methods. Data of all three methods is presented on FRT maps and it corresponds to tissue locations: i) tissue TFV concentrations measured by LC-MS/MS (squares), ii) proviral DNA detected by nested PCR (+/-), and iii) number of transduced cells (next to the white arrows pointing toward ovaries) identified by fluorescence microscopy.

## Discussion

In this work we studied the viral events and drug levels at 72-hours after exposure to a high dose of a non-replicative SHIV-LICh vector to explore how the TDF- IVR may protect from the earliest events after mucosal transmission. Our SHIV-LICh system allowed us to detect the initial sites of transduction within the macaque FRT by three different methods: A) a luciferase-based method via IVIS that allowed a rapid scan of the entire reproductive tract; B) a fluorescence microscopy-based method that allowed identification and characterization of single transfected cells, and C) a highly-sensitive nested PCR method that facilitated the detection of transduction events (proviral DNA) in the single tissue pieces across all animals. Interestingly, using these methods, we detect transduction events in the upper (uterus, ovaries, fallopian tubes) and middle (endocervix, upper vagina) reproductive tract even in TDF-IVR administered macaques.

TFV concentrations in the FRT were variable and were the highest in the tissues proximal to the ring. Lower drug levels were detected in distal tissues including ovaries. These results suggest that the upper reproductive tract is a potential site for initial infection despite the presence of the TDF-IVR.

Robust luciferase activity was identified in ovaries, fallopian tubes, and lymph nodes in seven out of ten IVR-treated animals, while very low luciferase activity was found in the cervix and vagina. While the IVIS method provides a survey of transduction events in the FRT, early viral events can only be detected on the surface of the tissue if multiple cells were transfected in a cluster to produce enough bioluminescence signal [2]. For that reason, a diffuse pattern of single viral transduction cannot be detected using IVIS. Therefore, further molecular analyses were critical to determine viral transduction in tissue with low or no luciferase activity detection.

Luciferase and mCherry expressing cells were found in the ovaries of five out of six IVR treated animals from the preclinical study as well as in the non-treated control animal (BB529). Spectral profiles of both reporter proteins demonstrated specificity of cells transduced with JRFL pseudotyped SIV virus. This result demonstrates the importance of the fluorescence microscopy method to confirm early viral events by finding transduced cells, since luciferase activity assay does not reveal single viral transduction events. Phenotyping analyses of transduced cells showed expression of CD4+ receptors in some of the mCherry and luciferase positive cells. This is in accordance with previously published data showing that CD4+ cells are primary targets during early infection in FRT (22–24). Furthermore, ovaries of the animal BB548 demonstrated positive luciferase activity, though no transduced cells were detected in these surveyed ovarian sections of this animal. To increase the likelihood of finding cells, additional tissue sections should be surveyed for the presence of transduced cells. It is very likely that some infection sites may be located at the other sites of the ovaries that were not necessarily included in the sectioning and were therefore not analyzed. Also, high auto fluorescence within ovarian tissue may have prevented us from identifying more specific transduced cells.

Measurements of the TFV levels throughout the FRT show variation with the highest levels of TFV accumulating in the upper vagina (maximum 6500 ng TFV/g tissue) and cervix (maximum 10,000 ng TFV/g tissue) near the site of the ring. The maximum TFV levels in the lower vagina (maximum 2600 ng TFV/g tissue), uterus (250 ng TFV/g tissue) and ovaries (400 ng TFV/g tissue) are statistically significantly lower compared to vagina and cervix. Our data suggest the possibility that the ovaries are more permissive to earliest infection events due to the lower levels of TFV accumulated in these distal tissue sites. Moreover, the mucus drainage gravitating from the upper toward the lower reproductive tract probably contributes to decreasing drug levels in the ovaries. The PCR analysis revealed that the entire genital tract except for the uterus was susceptible to transduction events. In addition to the FRT, proviral DNA was also detected in the lymph nodes of an animal (BB963), which is consistent with luciferase activity detected in this same tissue. Luciferase activity was detected also in the lymph nodes of several other animals (BB432, BB588, BB966 and 96P047) examined in the pilot study, but the presence of transduction events in these tissues was not confirmed by nested PCR. Similar findings reporting detection of early transduction events in lymph nodes of inoculated rhesus macaques were documented in our previous studies (17). Despite the highest drug levels detected around the endocervix and upper vaginal area where the ring was located, transduction events detected by nested PCR were identified in these areas in five of the six animals from the preclinical study. This suggests that achieving higher drug levels in the tissue does not necessarily provide protection from a high dose of SHIV. Susceptibility of the vaginal vault to infection may be due to differences in the intracellular environment that is tissue specific. Levels of dATP and dCTP were shown to be significantly higher in vaginal and cervical tissues compared to rectal tissue (25). Based on these findings, higher concentrations of exogenous nucleoside analogues such as TFV may be needed to successfully compete with cellular nucleotides for binding to the viral reverse transcriptase.

A limitation of this study is that the use of a single round infectious virus does not allow us to examine further spread of the virus within the tissue and into the bloodstream, a hallmark of a productive versus an abortive infection. We hypothesize that newly recruited immune cells in the FRT may not have accumulated enough drug to inhibit reverse transcription and protect them from SIV-based vector transduction. Moreover, migrating cells are by definition highly metabolically active with a high concentration of dATP and will require more TFV-dp to inhibit virus replication (25, 26). On the other hand, surrounding immune cells that were present in the FRT for a longer period and had enough time to accumulate higher drug levels may be better protected from viral transduction, and might prevent the viral spread outside the FRT that could result in systemic infection.

In order to test this hypothesis, further investigations utilizing replicative viruses mixed with single round infectious reporter viruses should help us to understand whether the use of the TDF-IVR ring could prevent viral spread from the first infected cells to other target cells as revealed by establishment of systemic infection. While this TDF-IVR was shown to protect macaques from infection from multiple low-dose SHIV challenge (13, 14), we observe incomplete protection from infection by the TDF-IVR as detected by vaginal challenge with our replication defective dual-reporter vector. Here we show that early sites of infection are detectable in TDF-IVR administered macaques after a single high dose viral challenge with a replication defective dual-reporter vector. The need for a single high-dose challenge with the dual-reporter required for signal detection should be considered in evaluating the outcomes in this model system. It has been shown in some nonhuman primates challenge studies that a high viral titer inoculation could overcome a PrEP formulation, while animals challenged with low viral dose remained protected (27) (28). A long-acting integrase inhibitor, cabotegravir, was demonstrated to efficiently protect macaques from repeated low-dose SHIV challenges (27). However, cabotegravir did not completely protect from a high viral dose challenge (28). Likewise, a long-acting injectable formulation of rilpivirine was also shown to have decreased efficacy against high viral dose (29). Furthermore, initial occult infections may be established without detectable viremia or immune responses in human populations. Studies of an early viral reservoir were shown to result in systemic viremia after discontinuation of fully suppressive therapy by ART (30). But the studies of in vivo PK/PD utilizing this system are not intended to provide insights into the prevention of systemic infection determined by the detection of viremia. Alternatively, this approach provides insights into topical PrEP function at the portal of transmission revealing potential deficiencies, such as the asymmetric distribution of the drug delivered in the vaginal vault by the TDF-IVR. Further studies will help us to understand the mechanism of protection by comparing topical drug delivery by TDF-IVR and orally administered TDF in high-challenge model.

Herein, we present a novel system that allows a comprehensive in vivo PK/PD study of topical PrEP formulations where drug levels and viral inhibition can be directly evaluated. In the current study, the function of a TDF-IVR was evaluated in the PTM vaginal challenge model with a single high viral dose of a non-replicative SIV based dual- reporter vector system that allows identification and analysis of potential target cells of the challenge inoculum. Transduction modeling initial viral events were found in, but not limited to the ovaries, cervix and vagina as expected based on previous studies (17). TFV levels were variable throughout the FRT in gradients of reducing tissue concentrations emanating from the IVR location. The drug released by the IVR in the vaginal compartment was able to reach the upper reproductive tract including the ovaries, although lower drug levels were detected. This observation suggests that IVR delivery systems may not protect the upper FRT as efficiently as the vaginal compartment. The highest drug concentrations were found proximal to the ring location; in the ectocervix and upper vagina compartments of the FRT. The detection of some transduction events overlapping with regions of high tissue drug levels reveal that viral reverse transcription was not completely inhibited with a high dose viral challenge despite the highest levels of TFV accumulated in these tissues. This is likely a case of the use of a single drug in this IVR formulation that competes with endogenous deoxyadenosine for inhibitory function. Unfortunately, it is not reasonable to address this issue by increasing the TDF concentrations. As shown in the early termination of Phase Ib clinical study of this TDF-IVR, ulcerations and significant increases in the levels of multiple inflammatory cytokines and chemokines in cervicovaginal fluid of sexually- active women administered with TDF-IVR compared to placebo group revealed drug release would need to be reduced for safety reasons (16). The high concentrations of TFV near the ring may be causing the observed adverse responses. Based on the insights gained in this initial study, this in vivo PK/PD approach can contribute important information to facilitate the development of effective topical PrEP formulations and maybe even provide insights into why oral PrEP that broadly disseminates drugs may be advantageous to topical PrEP in protecting high risk populations (1, 31).

## Materials and Methods

### Ethics statement

All animal experiments were conducted in accordance with protocols approved by Northwestern University and the Centers for Disease Control and Prevention Institutional Animal Care and Use Committee. This study was conducted with the recommendations in the Guide for the Care and Use of Laboratory Animals of the National Institute of Health (NIH).

### TDF-IVR study in non-human primates challenged with high viral dose

Six pigtail macaques were used for the pilot study and seven pigtail macaques for the preclinical study. All animals were given a single dose of 30 mg Depo-Provera to synchronize animals in their cycles. Four days later, TDF-IVR were administered to four animals in pilot study (BB432, BB588, BB401 and BB925) and six animals (BB535, BB963, BB548, BB187, 1.6348 and 1.8678) in the preclinical study. Two control animals (96P047 and BB966) in the pilot study and one animal (BB529) in preclinical study did not receive the TDF-IVR. All study animals were vaginally challenged with a high dose of single round non-replicating SIV-based reporter virus (TCID50 range: 10^5^-10^6^) twenty-five days (preclinical study) and twenty-eight days (pilot study) after IVR insertion. After viral inoculation animals were euthanized at 48 hours (animals in pilot study) and 72 hours (animals in preclinical study). A timeline for both studies is shown in Figure 1.

At necropsy, intact genital tracts and regional lymph nodes were isolated, stored in RPMI and shipped on wet ice overnight for further processing. The next morning tissue was analyzed for luciferase reporter expression. Whole FRT and lymph nodes were rinsed in PBS and scanned in the IVIS device to determine the background luminescence. Then the tissue was soaked in 100 mM d-Luciferin (Biosynth), while ovaries were injected with 100 mM d-Luciferin. After 10-15minutes at room temperature all tissue was examined for luciferase expression using IVIS. The genital tract was further divided into 5 regions: lower vagina, upper vagina/fornix, cervix, uterus and ovaries including fallopian tubes, and reimaged. Every region of the tissue was then dissected into smaller pieces and scanned again. Tissue sections with positive luciferase activity signal were further dissected into smaller bites for the third screen. Live Imaging Software was used for all analyses of luminescent signals. All dissected tissue was embedded in optimal cutting temperature (OCT) media (VWR) and frozen at -80 degrees Celsius. For the preclinical study, ovaries were cryosectioned and screened for the presence of transduced cells using fluorescence microscopy, while all other dissected domains of the FRT were used for nested PCR analyzes.

### SHIV-based dual reporter vector and viral production

In this study we apply a dual reporter genome that allows us to localize and phenotype cells transduced with a non-replicating SIV-based vector. Vector design was described previously (17). Briefly, the dual reporter genome carries firefly luciferase (18) and fluorescent mCherry (19) genes that are driven by a CMV promoter (Supplemental Figure 1). IRES (internal ribosome entry site) enables efficient expression of the mCherry gene whereas WPRE (Woodchucks Hepatitis Virus posttranscriptional regulatory element) at the 3’ end of the genome enables increased gene expression of the vector (20, 21). The 5’ end long terminal repeat (LTR) contains the SIV promoter for efficient virus production. The 3’ end LTR site has a self-inactivating mutation. The reporter genome does not contain any other viral genes, and is delivered by an SIV-based lentiviral vector.

An SIV-based pseudovirus vector was driven from the SIV3+ (SIVmac251) as described in (32). To generate pseudotyped reporter virus the 293T cells were transfected with 4 plasmids mixed with Polyethylenimine (PEI, Polysciences): dual reporter genome vector, SIV3+ packaging vector, JRFL envelope encoding plasmid and REV expression plasmid DM121. 48 hours post transfection supernatant containing pseudotyped virus was collected and filtered through 0.45 µm-sized pore filters. Virus was concentrated using a 20% sucrose cushion, followed by titration and infectivity assay (TCID50) on TZM-bl cells as previously described in (33) and stored at -80 ͦC. TCID50 ranged from 10^5^-10^6^ virions mL^-1^.

To identify vector-transduced cells that express mCherry and luciferase reporter genes frozen ovaries were cryosectioned and immunostained for fluorescence imaging analyses. mCherry was identified based on its auto fluorescent signal while the luciferase protein was immuno-stained with anti-firefly luciferase antibody (Abcam) labeled with Zenon AF647 Mouse Labeling Kit (Life Technology). Positive transduced cells were identified by detecting high fluorescent signals in both mCherry and CY5 (identifying AF647 labeled antibody for luciferase protein) channels. In addition, we employed a further criterion using the TRITC filter in an orange wavelength to characterize transduced cells. Dim emission in TRITC excludes detection of a broad-spectrum auto fluorescence of the tissue indicating specificity of mCherry auto fluorescence.

This viral reporter system was designed for a single round cell entry, viral transcription and proviral DNA integration. These reporter virions are therefore non-replicating viruses. This strategy allows us to localize early viral events based on the expression of reporter genes.

### Drug levels during the pilot and preclinical studies

Vaginal fluid samples were collected and assayed as previously described (13). Vaginal pinch biopsies (3-5mm) were collected on days 3 and/or 14/15 for drug level analysis. A section of vaginal tissues from the FRTs of the first group of macaques was collected at necropsy and processed by mechanical shearing (34) to isolate leukocytes for drug analysis at the cellular level. Concentrations of TFV and TFV-DP in vaginal tissues were determined by LC-MS as previously described (13).

### Drug PK post necropsy

TFV levels in tissue were measured by LC-MS/MS methods. The LLOQ for TFV was 50 ng TFV/g tissue. Tissue throughout the female reproductive tract was collected and sectioned to 100 mg pieces prior to the luciferin treatment. A solution of cold 50:50 acetonitrile:water spiked with 100 ng/mL ^13^C labeled TFV internal standard (Moravek, Brea, California) was added to the tissue with one 5 mm stainless steel homogenizing bead. Samples were homogenized for 10 min at 30 Hz and centrifuged 4 min at 10,000 rpm. Supernatants were filtered with 0.2 µm Nylon filters and evaporated to dryness on a vacuum centrifuge. Samples were reconstituted in water and injected on an HPLC- MS/MS system. Chromatographic separation was achieved using an Agilent Zorbax Eclipse XDB 2.1×150mm 5µ with mobile phases of 0.1% formic acid in water and 0.1% formic acid in acetonitrile at 0.3 mL/min with an Agilent 1200 HPLC. Quantification was achieved using a Bruker AmaZon X Ion Trap with Bruker Compass DataAnalysis QuantAnalysis version 2.1 software. The assay was linear in the range of 50-5,000 ng/g tissue.

### Immunostaining and fluorescence microscopy

Sixteen micron thick tissue cryosections were fixed on glass slides with a PIPES- formaldehyde mix (0.1 M PIPES buffer, pH 6.8 and 3.7 % formaldehyde) and washed with 1x PBS. Tissue was then blocked with 10% normal donkey serum/0.1% Triton X- 100/0.01% NaN3 followed by staining with Alexa Fluor 488 conjugated anti-human CD4 OKT 4) antibody (Biolegend) at 4° Celsius overnight (1:200 diluted in PBS). The next morning tissues were rinsed in 1X PBS and stained with rabbit polyclonal anti-firefly luciferase (Abcam) antibody, pre-labeled with Zenon AlexaFluor -647 mouse labeling kit according to manufacturer (Life Technology). DAPI was used to stain nuclei.

Immunostained cryosections were mounted with fluorescent mounting medium (DAKO) and covered with coverslips. Imaging was conducted with a DeltaVision inverted microscope. Transduced cells were visualized with a 60X objective lens using 2×2 paneled fields. Thirty z-scan stack images were acquired in 5 channels: DAPI, FITC (AlexaFluor 488), TRITC, mCherry and CY5 (AlexaFluor647), and deconvolved using softWoRx software (Applied Precision). Expression of mCherry and luciferase conjugated to AlexaFluor647 were analyzed by spectral imaging using Nikon AIR Laser scanning confocal microscope and Nikon Elements Software.

### Primer efficiency assay and nested PCR

293T cells (American Type Culture Collection) were grown in Dulbecco’s modified Eagle’s medium (Mediatech) supplemented with 10% fetal bovine serum, 100 Um L^-1^ penicillin, 100 µg mL^-1^ streptomycin and 292 µg mL^-1^ l-glutamine (Gibco). At 50% confluence cells were transduced with 1000 TCID50 JRFL pseudotyped vector in 10-cm plates for 24 hours followed by changing the media. Forty-eight hours post transduction cells were trypsinized, rinsed and collected for selection based on mCherry expression using a Beckman Coulter MoFlo system.

Genomic DNA was isolated from wild type (mCherry negative) and mCherry positive cells using Qiagen DNeasy Blood & Tissue Kit (Qiagen N.V.). DNA concentration was measured and calculations were made to prepare a genomic DNA mixture representing approximately 100 transduced cells and 3.79×10^4^ wild type cells in a total of 250 ng DNA corresponding to approximately 3.8×10^4^ cells. The DNA mixture was further tittered with genomic DNA of wild type cells and used in a primer efficiency assay to establish the sensitivity of nested PCR as described below.

Approximately three 40micron thick tissue cryosections were used to extract genomic DNA using above mentioned Qiagen DNeasy Blood & Tissue Kit. Tissue was analyzed for the presence of mCherry gene in a total of 250 ng DNA per reaction. DreamTaq polymerase (Promega) and primers were mixed with template DNA. Two sets of primers were used in this assay. To target the mCherry gene we used IRES and WPRE outer primers (IRES forward: 5’-ACATGTGTTTAGTCGAGG-3’ and WPRE reverse: 5’- CAGTCAATCTTTCACAAATTTTGTAATCC -3’) and mCherry inner primers (forward: 5’-CCGACTACTTGAAGCTGTCCTT -3’ and reverse: 5’- GTCTTGACCTCAGCGTCGTAGT -3’). The LTR sites were utilized as a positive control in the primer efficiency assay only, however not for screening (1) the animals’ tissues. Animals were previously infected with the SHIV162p3, and therefore only the presence of mCherry proviral DNA is a valid indicator of transduction in the macaque samples. To amplify the LTR sites we used LTR outer (forward: 5’-GCCTGTCAGAGGAAGAGGTTAG-3’ and reverse: 5’-GCCTTCACTCAGCCGTACTC -3’) and LTR inner primers (forward: 5’- TGGCTGACAAGAGGGAAACTC -3’ and reverse: 5’-CTCCTTCAAGTCCCTGTTCG -3’). Supplemental Figure 1 shows primer design on the dual reporter vector.

In the first round, PCR outer primers were used to generate outer PCR products. The cycling parameters were: 95 °C for 1 min 30 sec followed by 1 cycle, 95 °C for 30 sec, 45 °C for 30 sec and 72 °C for 1 min 50 sec followed by 20 cycles, and 72 °C for 10 min followed by 1 cycle, and the final step 4 °C forever. In the second round of PCR, 2 µL of the first PCR products were amplified with inner primers to generate final PCR products. The cycling parameters were: 95 °C for 5 min followed by 1 cycle, 94 °C for 30 sec, 51 °C for 30 sec and 72 °C for 45 sec followed by 35 cycles, and 72 °C for 5 min followed by 1 cycle, and the final step 4 °C forever. The nested PCR was then performed on a Bio-Rad iCycler Thermal Cycler system (Bio-Rad Laboratories).

All final PCR reactions were examined on 2% agarose gel and PCR products were visualized by ethidium bromide staining. PCR products were cut out from the gel and DNA was extracted using Quiagen QIAquick Gel Extraction Kit and sequenced with the inner mCherry primers. We then tested sensitivity of the nested PCR method. The primer efficiency assay demonstrated that by using a combination of the IRES/WPRE and mCherry primer sets we were able to detect up to 3 DNA copies per PCR reaction in a total of 250 µg of DNA (Supplemental Figure 2). This corresponds to 3 transduced cells in the mix of the wild type and transduced cells. As a positive control, we designed primers targeting the LTR sites that enable us to identify less than a single copy proviral DNA in our nested PCR reaction (Supplemental Figure 1, 2). Amplification of LTR elements resulted in higher production of PCR products because of the two LTR elements on each site of the molecule. Furthermore, amplification of longer PCR products such as 1.5 kb long fragments between IRES and WPRE elements is less efficient than amplifying shorter DNA fragments such as 587 bp long outer LTR products. Despite the high sensitivity of the LTR detection we could not use this set of the LTR primers to survey the tissue of the pigtail macaques because the animals had been challenged with SHIV162p4 viral inoculum prior to the beginning of our study. Due to the similarities of the LTR sequences in the dual reporter SIV-based virus driven from SIVmac251 and the SHIV162p3 driven from SIVmac239, we couldn’t use the LTR primers to distinguish between the two SIV strains.

**Supplemental material** is available online only.

TEXT S1, DOCX file, 17 KB.

FIG S1, PDF file, 50 KB

FIG S2, PDF file, 4 MB

FIG S3A, PDF file, 122 KB

FIG S3B, PDF file, 226 KB

## Acknowledgments

This work made use of the Integrated Molecular Structure Education and Research facility, which has received support from the SHyNE Resource (NSF ECCS-2025633), the IIN, and Northwestern’s MRSEC program (NSF DMR-1720139). Spectral imaging work was performed at the Northwestern University Center for Advanced Microscopy generously supported by NCI CCSG P30 CA060553 awarded to the Robert H Lurie Comprehensive Cancer Center. We thank the Robert H. Lurie Comprehensive Cancer Center of Northwestern University in Chicago, IL, for the use of the Flow Cytometry Core Facility, which provided cell sorting service. The Lurie Cancer Center is supported in part by an NCI Cancer Center Support Grant #P30 CA060553. Professor Elena Martinelli from Northwestern University for providing her scientific feedback. We acknowledge the following members of the CDC DHAP Laboratory Branch/Preclinical Evaluation Team for their contributions to our nonhuman primate research: David Garber, James Mitchell, Ryan Johnson, Shanon Ellis and Kristen Kelley.

## Disclaimer

The findings and conclusions of this manuscript are those of the authors and do not necessarily represent the official views of the Centers for Disease Control and Prevention.

---

**Fig. S1.**
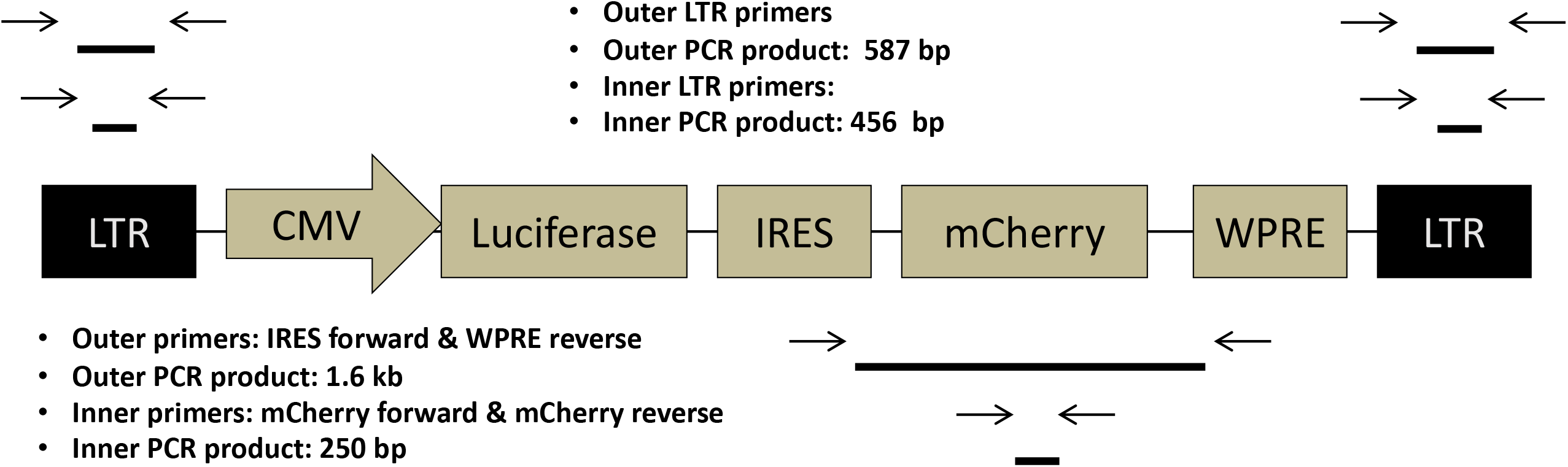
Scheme of dual reporter genome LICh and primer design for nested PCR.

**Fig. S2.**
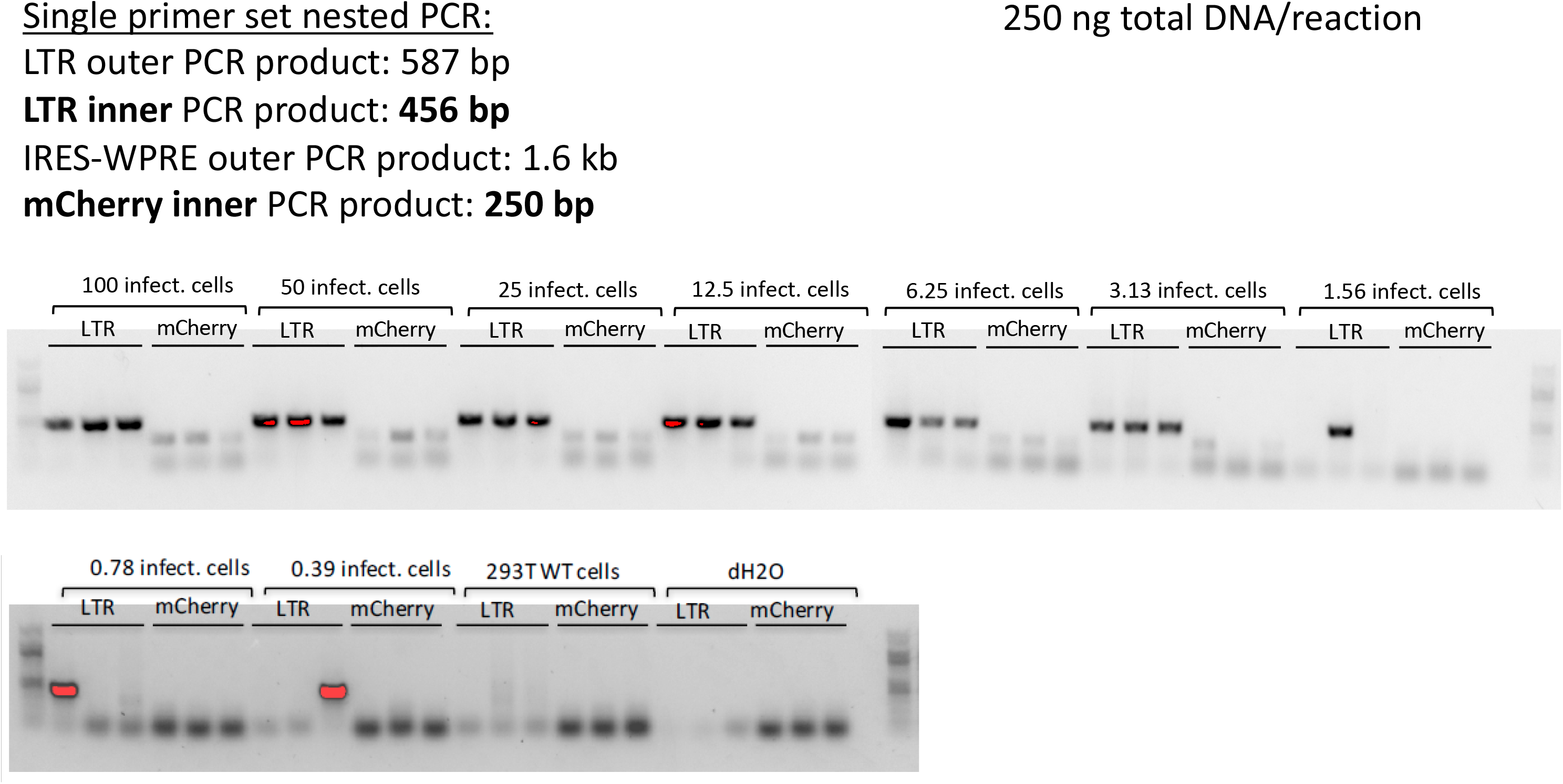
Primer efficiency assay. Genomic DNA mixture representing 100 infected cells in 250 ng DNA was further diluted and used in primer efficiency assay using IRES/WPRE and mCherry primer set and LTR primers as a positive control

**Fig. S3.**
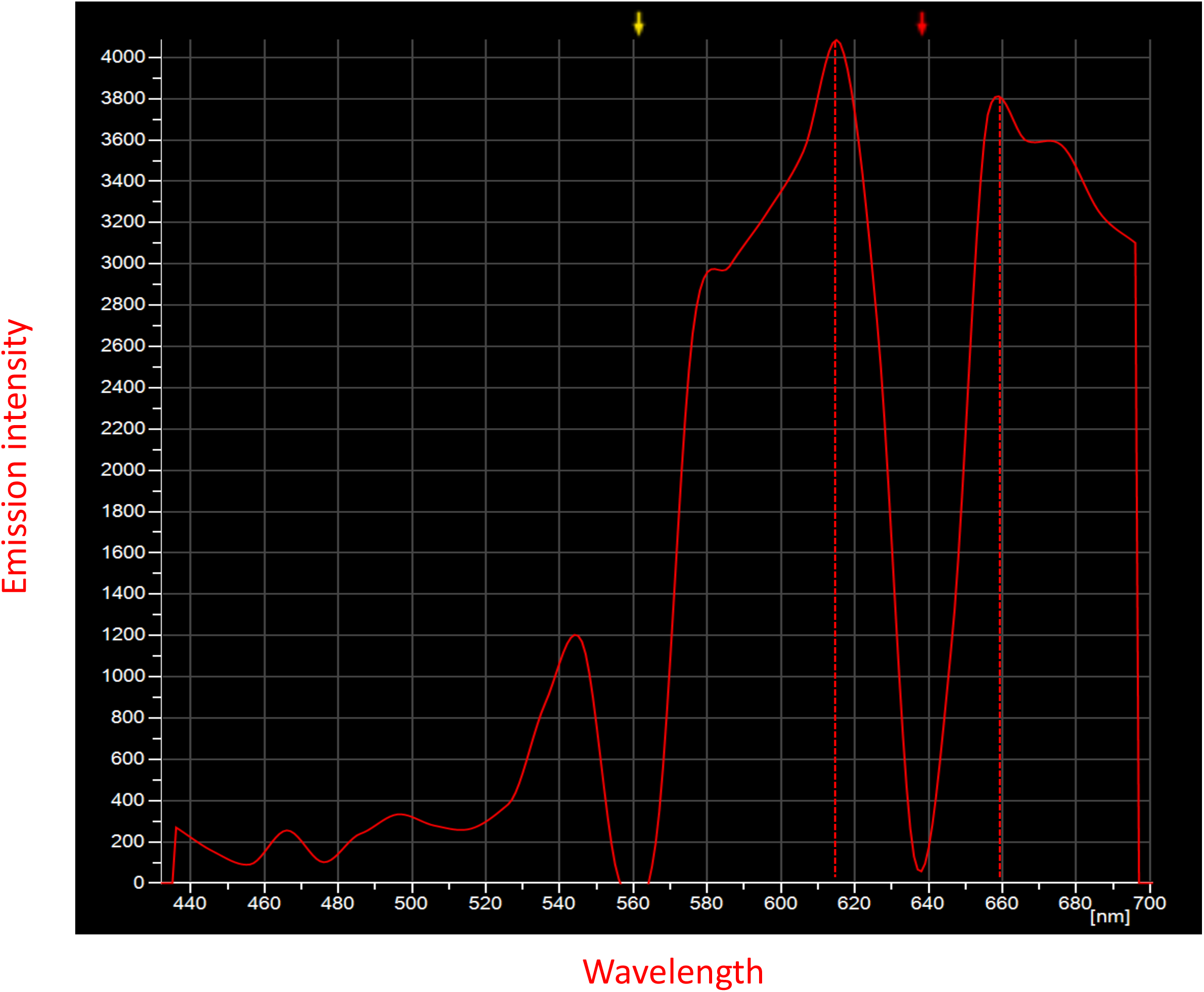
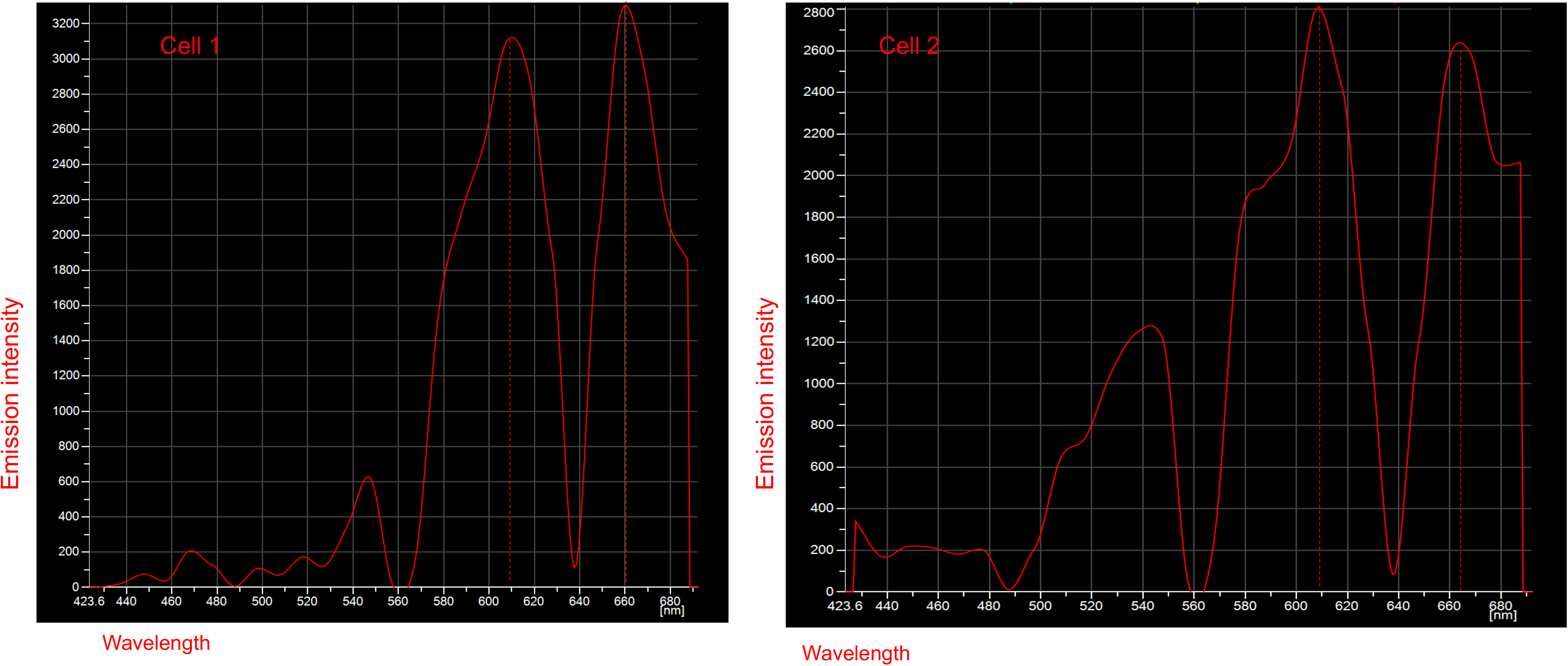
Spectral profile analyzes of transduced cells. Spectral emission profile matching the defined profile of mCherry (610 nm) and luciferase seen in CY5 (665 nm) of ovarian tissue of TDF-IVR-treated animal BB535 (3A) and BB548 (3B).

